# The genetics of transcriptional responses to stress in yeast

**DOI:** 10.64898/2026.05.29.728873

**Authors:** Noah Alexander, James Boocock, Heriberto Marquez, Joshua S Bloom, Leonid Kruglyak

## Abstract

Alteration of gene expression levels is a key cellular strategy for responding to internal and external challenges. While expression quantitative trait loci (eQTLs) are known to influence steady-state transcript levels, their role in mediating rapid physiological transitions remains poorly understood. Using a high-throughput, “one-pot” single-cell RNA-seq approach, we mapped eQTLs in yeast immediately before and after two acute environmental shifts: salt perturbation of actively cycling cells and nutrient repletion of cells after one week of starvation. We defined and characterized major cell populations in different physiological states and identified context-specific eQTLs as well as genetic determinants of state occupancy. We focused on distant eQTL hotspots, which capture the majority of expression heritability in yeast crosses. We found that the effects of hotspots on gene expression are highly dynamic in response to acute environmental perturbations, and that they influence biological processes directly relevant to adaptation to the new conditions. By integrating gene set enrichment analysis with computational prediction of variant effects, we prioritized candidate causal genes and variants underlying the hotspots. We observed extensive overlaps between hotspot loci, physiological state QTL, and fitness QTL. These findings reveal how regulatory variation rapidly modulates cellular responses to environmental changes, suggesting a mechanistic bridge between genetic variation, gene expression, and organismal fitness.

## Introduction

The genetics of gene regulation can provide considerable insights into the mechanistic basis of organismal trait variation^1–3^. Extensive mRNA expression quantitative trait locus (eQTL) mapping studies have been conducted over the last two decades. In humans, these studies have focused on the effects of common variants on expression of nearby genes (local eQTL)^4,5^. Studies in model organisms have revealed extensive effects of genetic loci on the expression of unlinked genes (distant eQTL)^6^. The effect sizes of individual distant eQTL are on average smaller than those of local eQTL, but in aggregate explain more of the overall expression variance in populations^7,8^. Distant eQTLs are more likely to mediate the effects of DNA sequence variation on cellular and organismal traits^7^. A key advantage of mapping studies in yeast is that they have high statistical power to identify both distant and local eQTL, as well as fitness traits^1,8–10^.

An important result from eQTL studies in model organisms is the existence of eQTL “hotspots”—loci that influence the expression levels of many more unlinked genes than would be expected by chance^6,8^. The affected transcripts are often functionally related, suggesting that hotspots result from genetic differences that alter the regulation of specific biological programs. Investigators can often identify candidate causal genes underlying eQTL hotspots by focusing on genes located within a hotspot that are functionally related to the transcripts affected by the hotspot. In some cases, there may be no known connection between a causal gene and the hotspot transcripts, and this can reveal new regulatory relationships. Most of the causal variants underlying hotspots in yeast have been detected in coding regions^1,8^. The effects of distant eQTLs, including those that result in hotspots, tend to be more dependent on the environment and on physiological state than those of local eQTLs^11–13^. To date, experimental constraints have limited our ability to conduct eQTL studies across many genetic backgrounds and environmental contexts.

Single-cell eQTL mapping can overcome these constraints. Unlike prior approaches that require separate cultures and bulk gene expression measurements for each individual or strain, single-cell RNA-seq enables ‘one-pot’ eQTL mapping, in which cells with different inherited combinations of genetic variants are pooled and profiled together in a single experiment, with each cell’s genotype inferred from its transcriptome^14–16^. This design allows multiple crosses and environments to be analyzed within the same study with lower cost and effort. In addition, this approach facilitates eQTL mapping over a time course following environmental perturbation, providing access to information about genetic effects on gene regulation that cannot be obtained from steady-state measurements alone. Single-cell RNA-seq data enables classification of the physiological state of each cell, allowing eQTL mapping within specific states and identification of genetic variants that influence occupancy of those states.

Here, we sought to extend previous studies by examining additional environmental conditions and by characterizing physiological states that yeast cells can occupy outside of conventional cell cycle phases. For instance, the set of states that yeast cells can occupy during nutrient starvation and after nutrient repletion has not been thoroughly characterized at the single-cell transcriptome level, although it is known that cultures grown in these conditions are highly heterogeneous with respect to cellular properties, including each cell’s potential to reenter the cell cycle^17–19^. We therefore carried out single-cell eQTL mapping in yeast cultures that entered stationary phase through gradual nutrient depletion and were then refed, as well as in exponential phase cultures that were acutely exposed to a high concentration of salt (Figure 1). We used the single-cell transcriptome data to assign physiological state classifications to each cell and looked for genetic determinants of the occupancy of each state. We repeated these studies in two well-studied yeast crosses: one between the lab strain BY4741 (BY) and the wine strain RM11 (RM), and one between the soil strain CBS2888 (CBS) and the clinical strain YJM981 (YJM). We combined these eQTL and state occupancy QTL mapping results with extensive fitness QTL data we previously generated in the same crosses to generate hypotheses about causal pathways through which genetic variation influences gene expression, physiological state, and fitness^10^.

**Figure 1.**
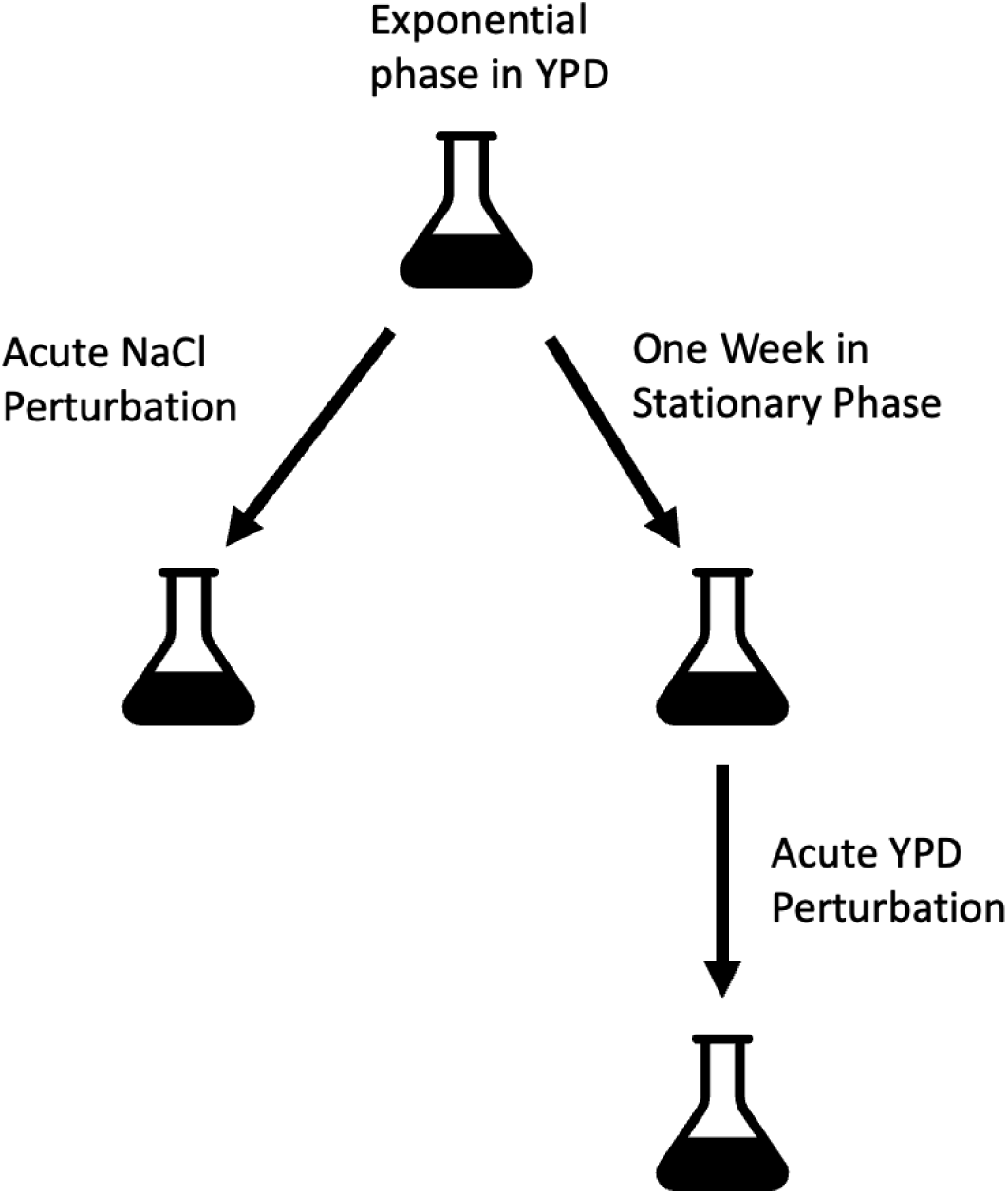
Schematic of the sampling scheme used in this study. The BYxRM cross was sampled under all conditions, whereas the CBSxYJM cross was only sampled across the salt perturbation time course (exponential phase in YPD and acute NaCl perturbation).

## Results

We carried out single-cell RNA-seq on segregant pools from the two crosses (BYxRM and CBSxYJM). For both crosses, we collected samples from mid-log phase cultures grown in rich medium (yeast extract peptone dextrose; YPD) and thirty minutes after a 0.7M salt perturbation (Figure 1; see methods). For the BYxRM cross, we also allowed a mid-log phase culture to reach stationary phase and remain in it for one week before refeeding the culture with fresh medium. We collected samples just prior to and ten minutes after refeeding (see methods). Between 10,000 and 20,000 cells were analyzed for each sample (Table 1).

**Table 1.** Single-cell transcriptome yields and data metrics for segregants in each condition.

| Cross/Condition | N cells | Mean reads/cell | Median genes/cell | Median nUMI/cell | Total genes detected |
| --- | --- | --- | --- | --- | --- |
| BYxRM YPD mid-log | 16,093 | 29,363 | 1,776 | 10,016 | 6,628 |
| BYxRM YPD mid-log + 30min NaCl | 14,652 | 36,425 | 1,904 | 9,384 | 6,682 |
| CBSxYJM YPD mid-log | 18,194 | 28,696 | 2,120 | 15,506 | 6,401 |
| CBSxYJM YPD mid-log + 30min NaCl | 18,026 | 25,035 | 1,870 | 10,864 | 6,429 |
| BYxRM SP 1wk | 10,472 | 36,752 | 498 | 1,215 | 6,377 |
| BYxRM SP 1wk + 10min YPD | 19,682 | 26,068 | 1,503 | 4,320 | 6,421 |

### Environmental stress response signatures in the segregant data

Yeast cells exposed to conditions such as salt stress and nutrient limitation are known to activate the environmental stress response (ESR)^20–22^. This response involves induction of a well-studied set of stress-related genes (iESR) as well as repression of ribosomal protein (RP) and ribosome biogenesis (RiBi) genes. In our data, we observed much higher iESR gene set activity in samples collected following acute salt perturbation and during stationary phase than in samples from mid-log phase or following nutrient repletion (Figure 2). The ribosomal protein gene set activity was low in stationary phase samples and much higher in the other conditions. We only observed appreciable ribosome biogenesis gene set activity in the mid-log phase cultures and in stationary phase cultures after nutrient repletion. These observations are consistent with the expectations that iESR activity should be higher in stressful conditions than during unimpeded proliferation, and that cells should expend resources to produce new ribosomes primarily in the presence of nutrients and in the absence of acute stresses^20,21^.

**Figure 2.**
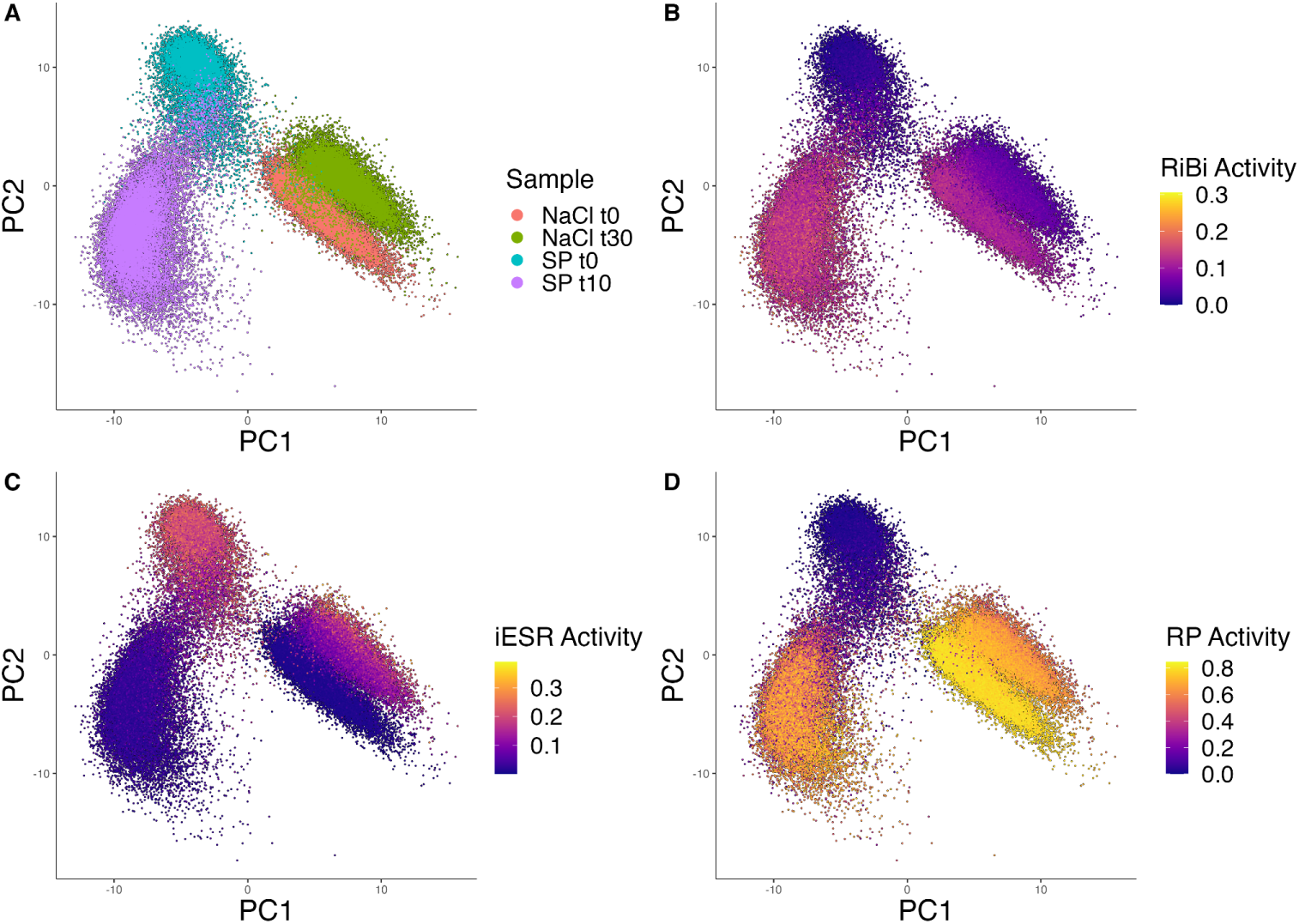
ESR gene set activity quantification in single-cell transcriptome profiles of segregants of the BYxRM cross. Panel A displays the experiment and timepoint that each single-cell transcriptome measurement derives from (“SP” denotes stationary phase) and panels B-D display continuous activity values for RiBi, iESR, and RP gene sets.

### Genetic mapping of ESR gene set activity traits

While the broad trends in ESR gene set activity across the sampled conditions corresponded to our expectations, we noticed that there were differences in gene set activity within each condition (Figure 2). To examine whether genetic variation contributes to these differences, we conducted linkage mapping of ESR gene set activity traits in each of the six samples (Figure 3). We treated iESR activity, RP activity and RiBi activity as distinct traits (see methods). These analyses revealed 44 QTL with LOD above 10 that were linked to at least one ESR gene set activity trait in at least one sample (Figure 3, see methods). Many of these QTL were observed for multiple ESR activity traits in multiple samples. This result is not surprising because the three stress-related gene sets are coordinately-regulated in cultures exposed to different stress conditions^20,21^. We note, however, that several QTL were specific to individual gene sets or time points, suggesting that the ESR is not entirely monolithic in its genetic control.

**Figure 3.**
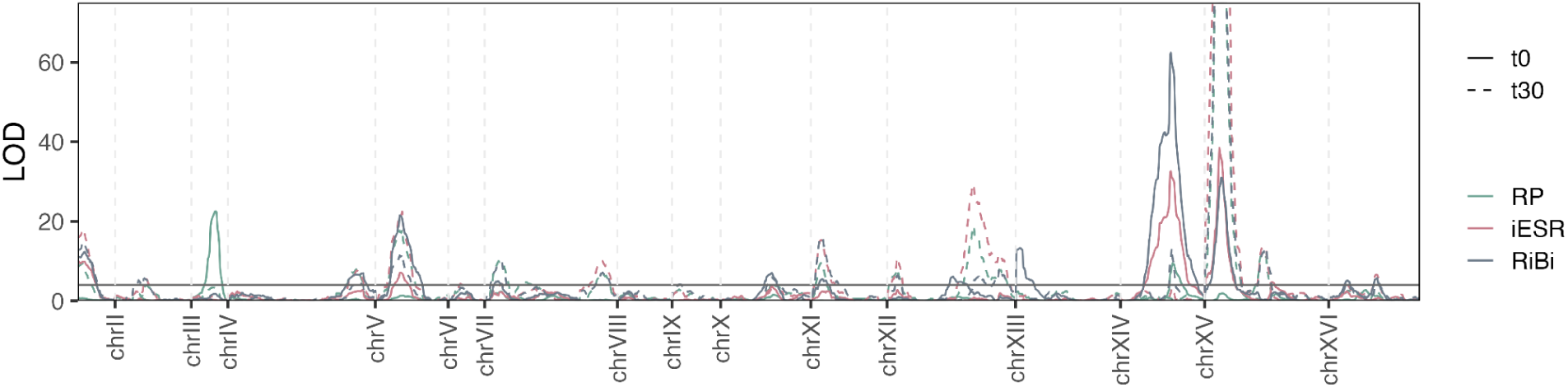
ESR gene set activity QTL (BYxRM salt experiment). Individual traces correspond to genetic mapping of iESR, RP, and RiBi gene set activity traits in the t0 (solid lines) and t30 (dashed lines) time points.

### Physiological state definition and cell classification

We used single-cell RNA-seq data to identify physiological states in each sample and to classify the state of each cell. States were defined as major populations of cells with distinct expression patterns by Leiden community detection on the top PCs of sctransform-normalized data, followed by manual annotation using known marker genes (see methods)^23^. We first focused on the salt perturbation experiments. In the BYxRM cross, we identified seven states in the mid-log phase cultures before salt stress and six after the perturbation. In the CBSxYJM cross, we identified six states in the mid-log phase cultures before salt stress and eight after the perturbation. Examination of expression patterns in these states showed that most corresponded to conventional cell cycle phases with an additional ‘stress’ state in the samples after salt stress, characterized by expression of several stress-induced genes^22^ (see methods). These results are consistent with prior single-cell work involving yeast cultures in mid-log phase and after an acute salt perturbation^14,22^.

Much less is known about the set of states that yeast cells occupy during stationary phase^17,18^. Bulk transcriptomics studies have shown that cells in stationary phase exhibit expression of genes related to mitochondrial function, beta oxidation of fatty acids, ATP generation, glyoxylate cycle, and multiple stress responses, but these studies could not identify subpopulations of cells contributing to these signatures^20,24^. We identified six distinct states in stationary phase yeast cultures after one week (Figure 4). Stationary phase cells are known to contain roughly 30-fold fewer mRNA molecules than cycling cells, and accordingly we observed lower UMI counts in these samples (Table 1)^17,18,25^. Two states are related to fatty acid metabolism and peroxisomal function, a pathway and organelle that are critical for energy generation in the absence of external nutrients. Two states are defined by the expression of stress-related genes (those related to misfolded protein response and those related to oxidative stress response). One state is distinguished by expression of salvage pathway genes. We also observed a population of cells expressing ribosomal protein transcripts but not ribosomal biogenesis transcripts, a pattern distinct from what would be expected in cells that are cycling or returning to the cell cycle ( Figure 5).

**Figure 4.**
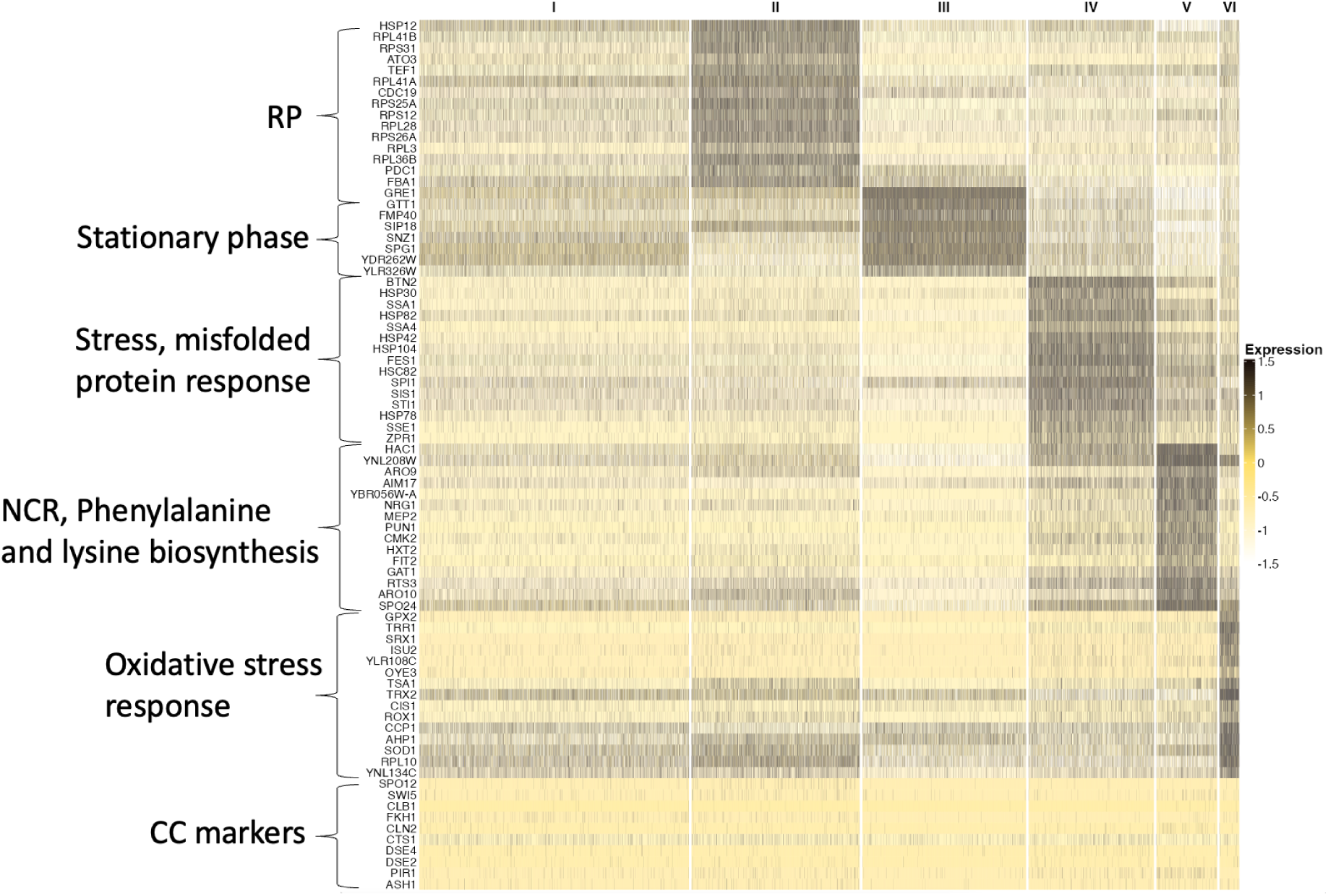
A heatmap of single-cell gene expression in BYxRM segregants after one week in stationary phase. Annotations to the left of the heatmap reflect functional categories of major blocks of expression. Cell cycle genes are visualized here to illustrate their low expression in the stationary phase context. RP and NCR denote ribosomal protein and nitrogen catabolite repression genes, respectively.

**Figure 5.**
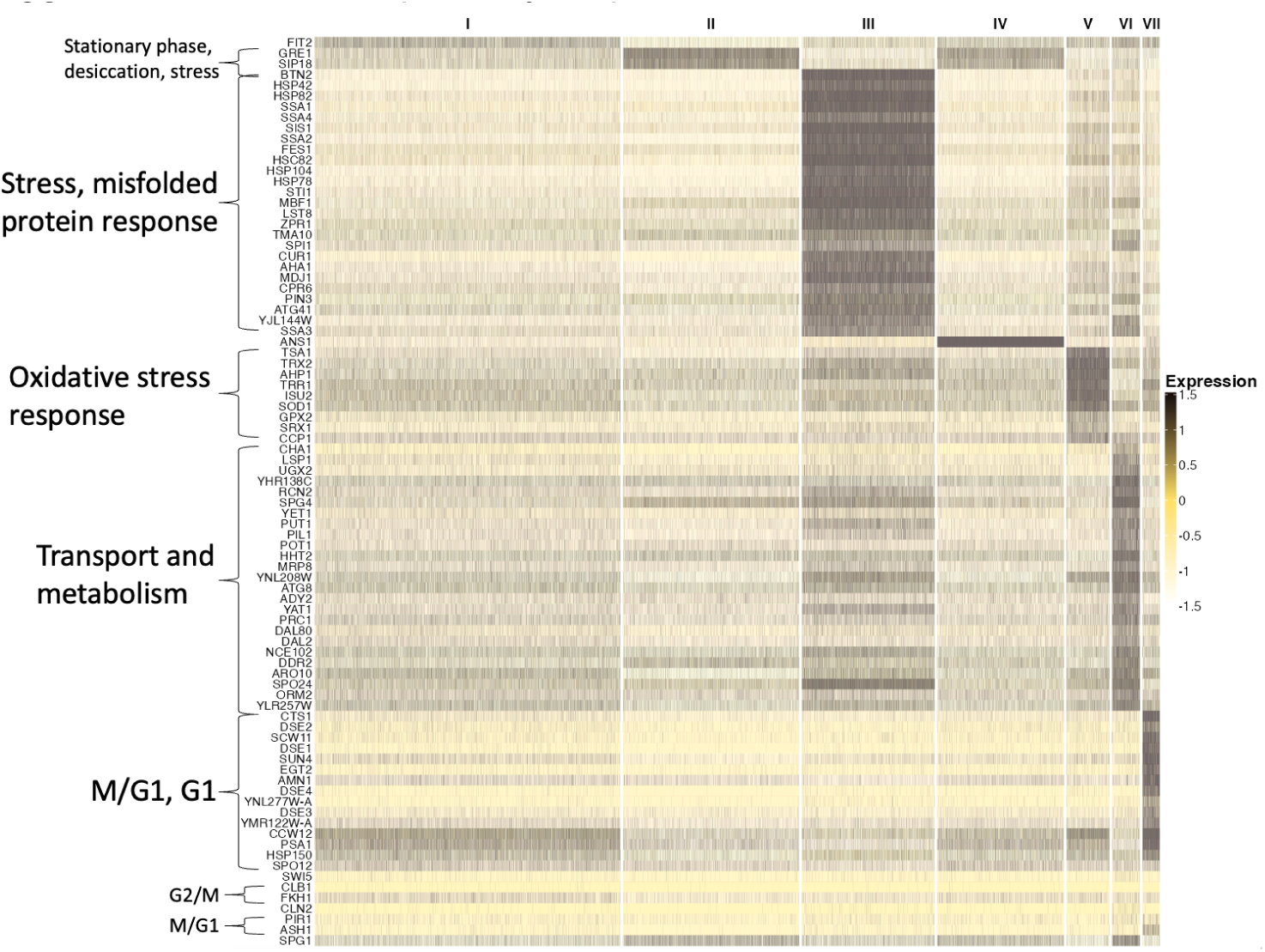
A heatmap of single-cell gene expression for BYxRM segregants ten minutes after a stationary phase culture was perturbed with rich medium. Annotations to the left of the heatmap reflect functional categories of major blocks of expression.

We next analyzed the sample collected 10 minutes after the addition of nutrients to a stationary phase culture. Extensive transcriptional shifts have been observed immediately after nutrient repletion of such cultures, and these shifts are thought to play a key role in determining whether an individual cell returns to the cell cycle^19,26,27^. Bulk studies have highlighted three transcriptional shifts: (1) rapid induction of transcripts related to ribosome biogenesis and translation, (2) a moderate spike in expression of nutrient transporters, cell-wall biosynthesis genes, sulfur metabolism genes, and other homeostatic regulators, and (3) gradual repression of genes involved in mitochondrial function, cellular respiration, and stress^21,28,29^. These changes are in line with the need of the cells to produce ribosomes to facilitate cell division, to import nutrients from the external environment, and to return from respiration to fermentation, the preferred mode of energy production. Ribosomal biogenesis gene expression is observed in our perturbed dataset and a small number of cells express early cell cycle markers (Figure 5). This suggests that these populations are preparing to enter the cell cycle and highlights the overall heterogeneity with which stationary phase cultures respond to nutrient stimulation. Two populations were distinguished by expression of unfolded protein response and oxidative stress response transcripts. Two populations exhibit expression of genes related to cell wall regulation, transporters, glycolysis, and biosynthesis of amino acids. One population exhibits expression of known stationary phase markers. The set of distinguishing markers for this population identified through differential expression analysis include genes involved with cytoplasmic translation (see methods). A final population also exhibits expression of *SPG1* and *GRE1* along with expression of *ANS1*, a putative GPI/membrane protein with loose connections to vacuolar function^30^. *ANS1* expression is very low elsewhere in the dataset. The markers for this cluster include genes related to cytoplasmic translation and rRNA processing. Together, the characteristics of this population suggest that it is transcriptionally responsive to the nutrient shift.

### Genetic mapping of physiological state occupancy traits

Shifting between physiological states in response to internal and external conditions is critical for cellular development and survival^19,20,31^. Although genetic factors are known to influence cell cycle phase occupancy in rapidly growing yeast cultures, similar analyses have not been extended to physiological states outside of the cell cycle^14^. To address this gap, we conducted genetic mapping of occupancy in each of the above-described physiological states (see methods).

We were especially interested in the genetic basis of variation in states observed during stationary phase and immediately following nutrient repletion. Although the genetics of quiescent state occupancy has been difficult to study, we presume one or more of our clusters corresponds to quiescence^17,18^. We detected substantially more QTL in the stationary phase experiment than in the salt perturbation experiment. In the BYxRM cross, we identified 16 QTL at stationary phase t0 and 12 at stationary phase t10, compared to one at NaCl t0 and four at NaCl t30 (Figure 6). Many of these loci also overlap eQTL hotspots, environmental stress response gene set activity QTL, and/or fitness QTL. Previously-identified eQTL hotspot loci (for example, those driven by variation in *IRA2* and *MKT1* in BYxRM and *CYR1* in CBSxYJM) influenced occupancy of many physiological states^8,14^. In addition, we identified several other loci that influence these traits. In several cases, occupancy in similar physiological states is influenced by the same loci. For example, a locus near the center of chrXV influences occupancy of misfolding stress-related clusters at both stationary phase-related time points (Figure 6).

**Figure 6.**
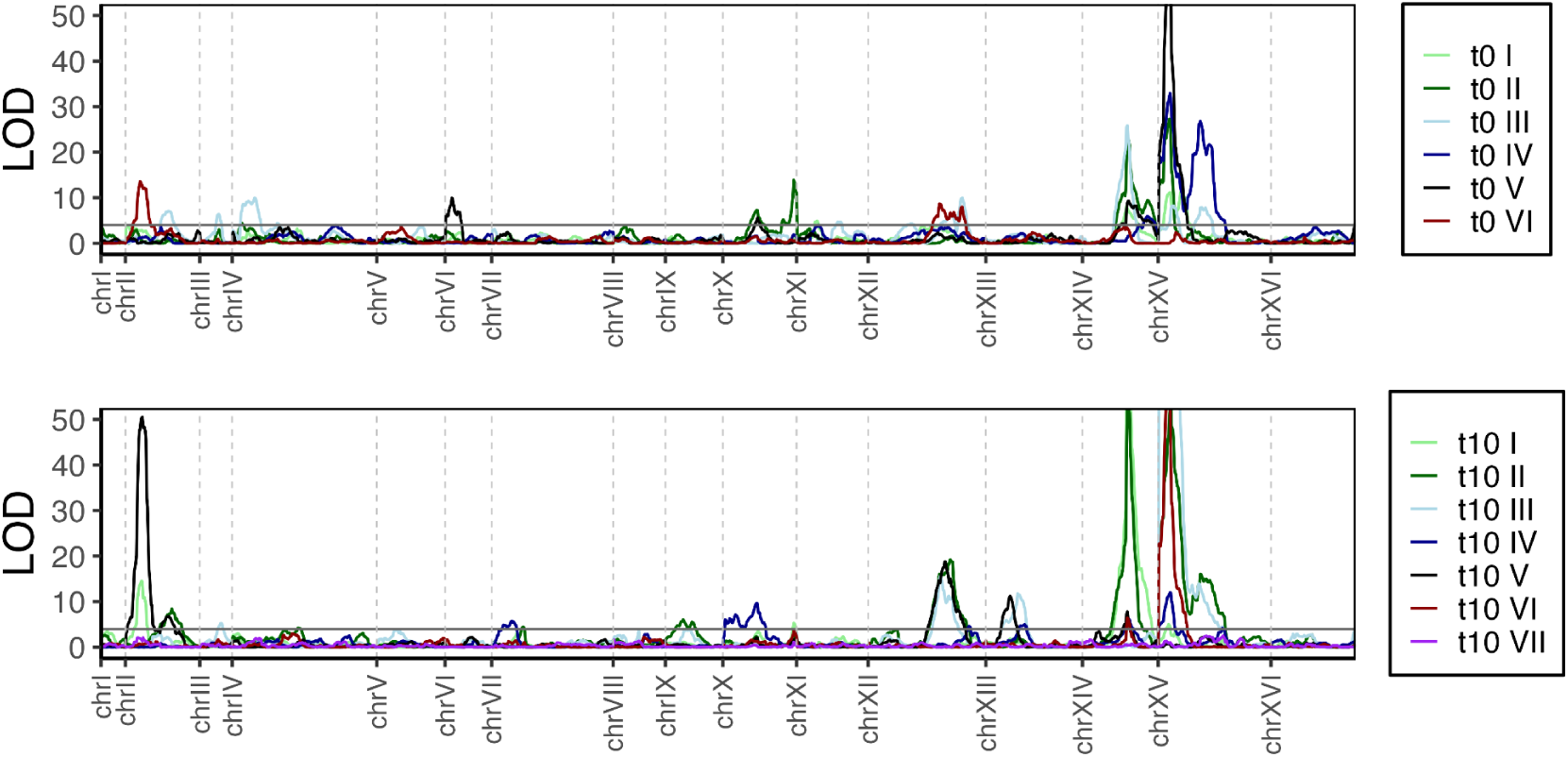
Overlaid LOD traces corresponding to genetic mapping of occupancy in each state identified in the stationary phase rich media perturbation experiment.

**Figure 7.**
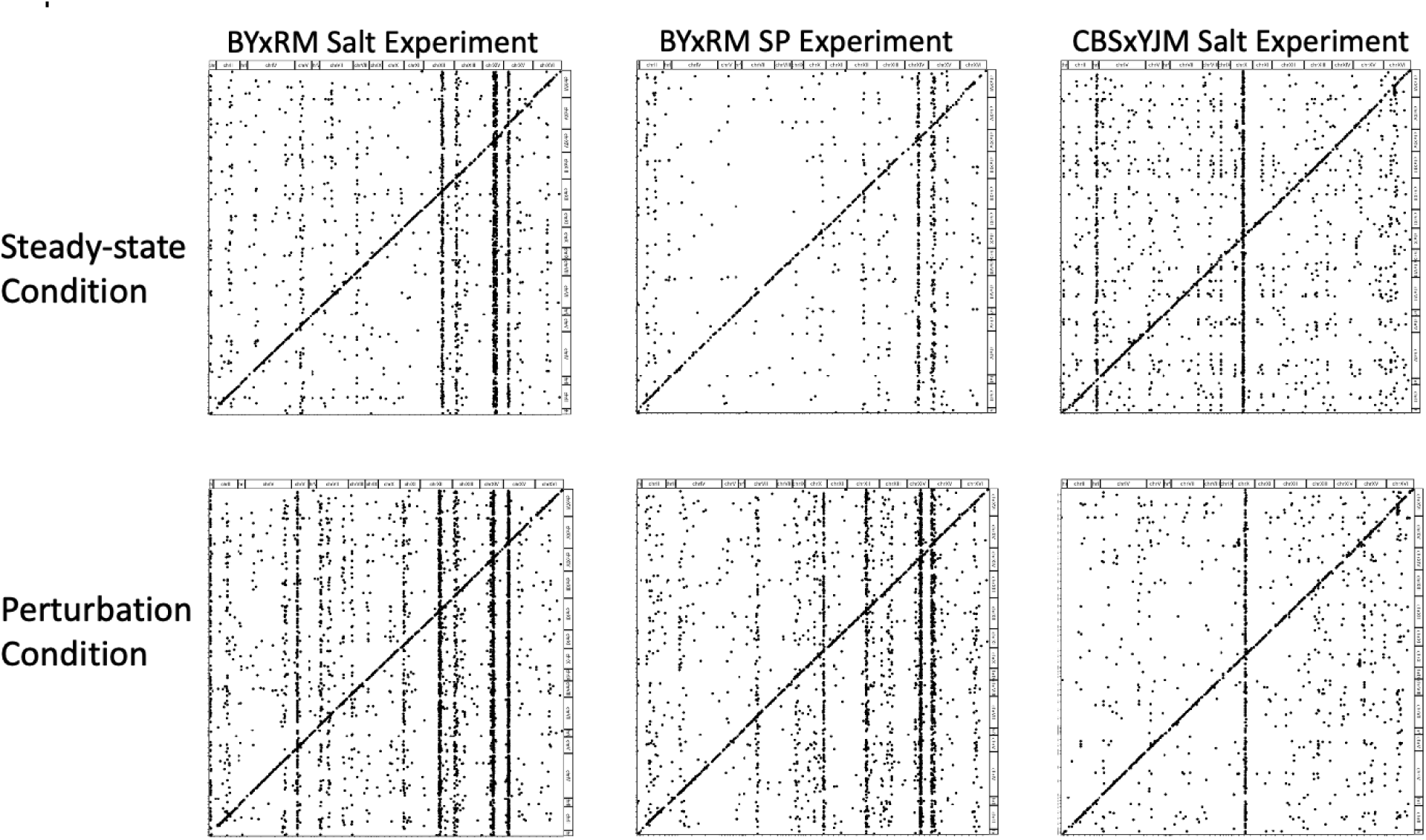
eQTL maps for each cross:condition combination. The rightmost column includes eQTL maps derived from the CBSxYJM cross and the leftmost two columns include maps from BYxRM salt and stationary phase experiments.

### eQTL mapping

We conducted separate eQTL mapping in each cross and sample with awareness of state assignments (see methods). At the combined level, we identified a total of 7,123 local eQTL and 16,646 distant eQTL across the study (Table 2). We used an FDR threshold of 5% for local and global scans.

**Table 2.** eQTL mapping results across local and global scans at the combined level. The numbers in parentheses in the hotspots column are the ranges of linking transcripts across hotspots in each condition.

| Cross/Condition | Local eQTL | Distant eQTL | eQTL Hotspots |
| --- | --- | --- | --- |
| BYxRM YPD mid-log | 1,146 | 2,610 | 9 (28-692) |
| BYxRM YPD mid-log + 30min NaCl | 1,348 | 5,546 | 11 (34-264) |
| CBSxYJM YPD mid-log | 1,075 | 2,255 | 21 (34-264) |
| CBSxYJM YPD mid-log + 30min NaCl | 1,358 | 1,605 | 7 (50-2042) |
| BYxRM SP 1wk | 840 | 1,064 | 8 (15-230) |
| BYxRM SP 1wk + 10min YPD | 1,356 | 3,566 | 11 (26-237) |

### Local eQTL

In the BYxRM salt perturbation experiment, we identified 1,778 local eQTL at a false-discovery rate (FDR) of 5%. Of these, 40.3% were detected both before and after salt stress, 24.2% were detected only before salt stress, and 35.5% were detected only after salt stress (Figure 8). In the CBSxYJM salt perturbation experiment, we identified 1,760 eQTL, with 38.2%, 38.9%, and 22.8% detected as above (Figure S21). In the BYxRM stationary phase experiment, we identified 1,739 eQTL, with 26.2% detected both in stationary phase and after nutrient repletion, 22% detected only in stationary phase, and 51.7% detected after nutrient repletion (Figure 8).

**Figure 8.**
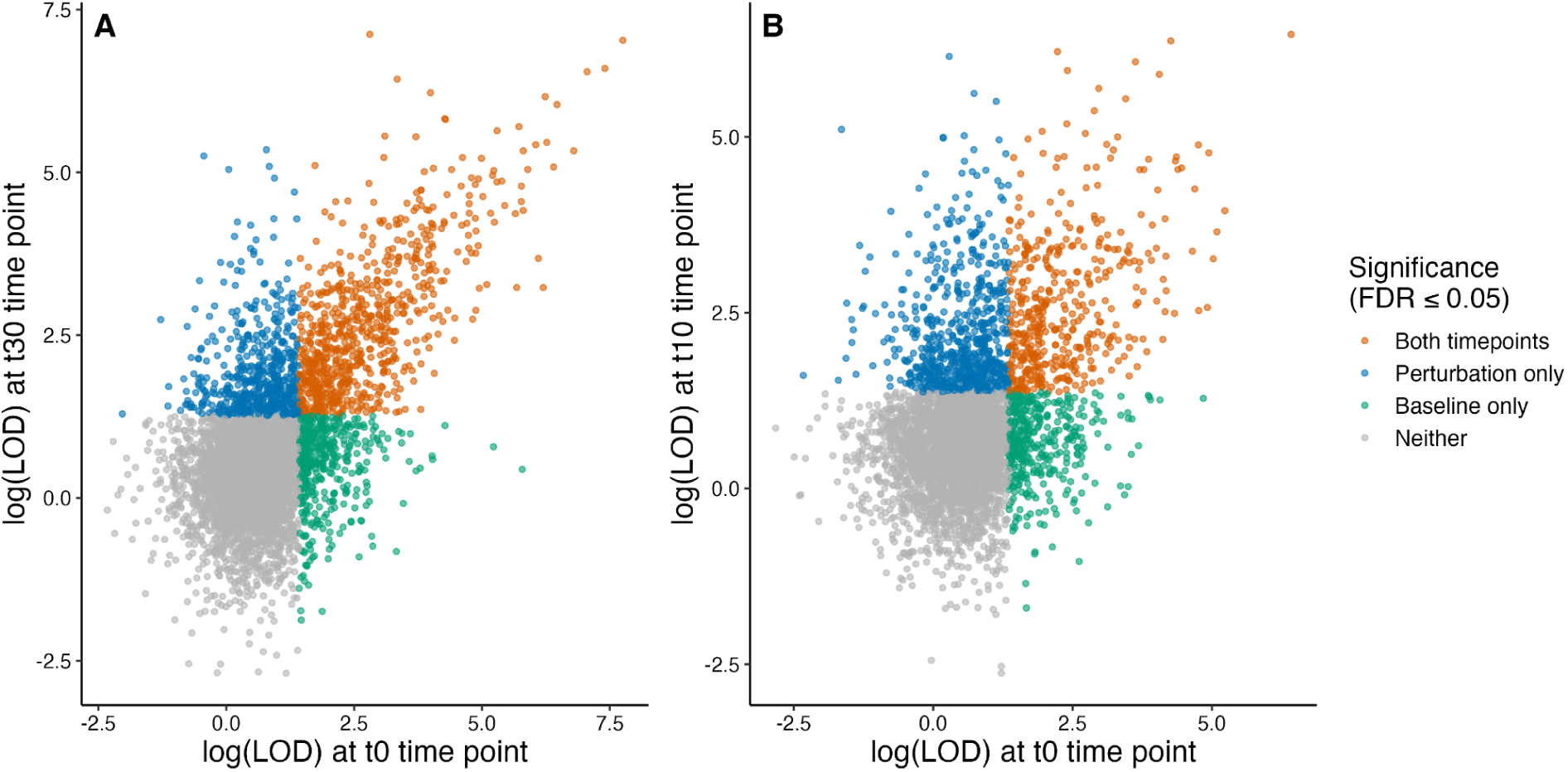
Local eQTLs mapped in the BYxRM perturbation experiment. Panel A shows LOD scores for transcripts corresponding to the NaCl t0 and NaCl t30 conditions. Panel B shows LOD scores for transcripts corresponding to the SP t0 and SP t10 conditions.

### eQTL hotspots

Distant eQTLs were significantly enriched at 44 unique hotspots—regions of the genome containing more distant eQTL than would be expected if they were randomly distributed throughout the genome (see methods, Figure 9)^6,8^. The functions of the transcripts linked to each hotspot were frequently closely related to the culture conditions in which the hotspots were detected, especially following salt stress and exit from stationary phase, as will be further discussed below. Most hotspots were detected in one or a small number of cell states (Figure 9). Consistent with prior work, several hotspots had allelic effects that changed the expression of all linked transcripts in the same direction^8,11^. For instance, of the 18 hotspots identified in the CBSxYJM salt perturbation experiment, 7 fell into this category (see SI Tables 1 and 2). Such hotspots reflect especially clear and interpretable genetic effects on gene regulation. Other hotspots had allelic effects that changed the expression of different linked transcripts in opposite directions, suggesting more complex regulatory scenarios.

**Figure 9.**
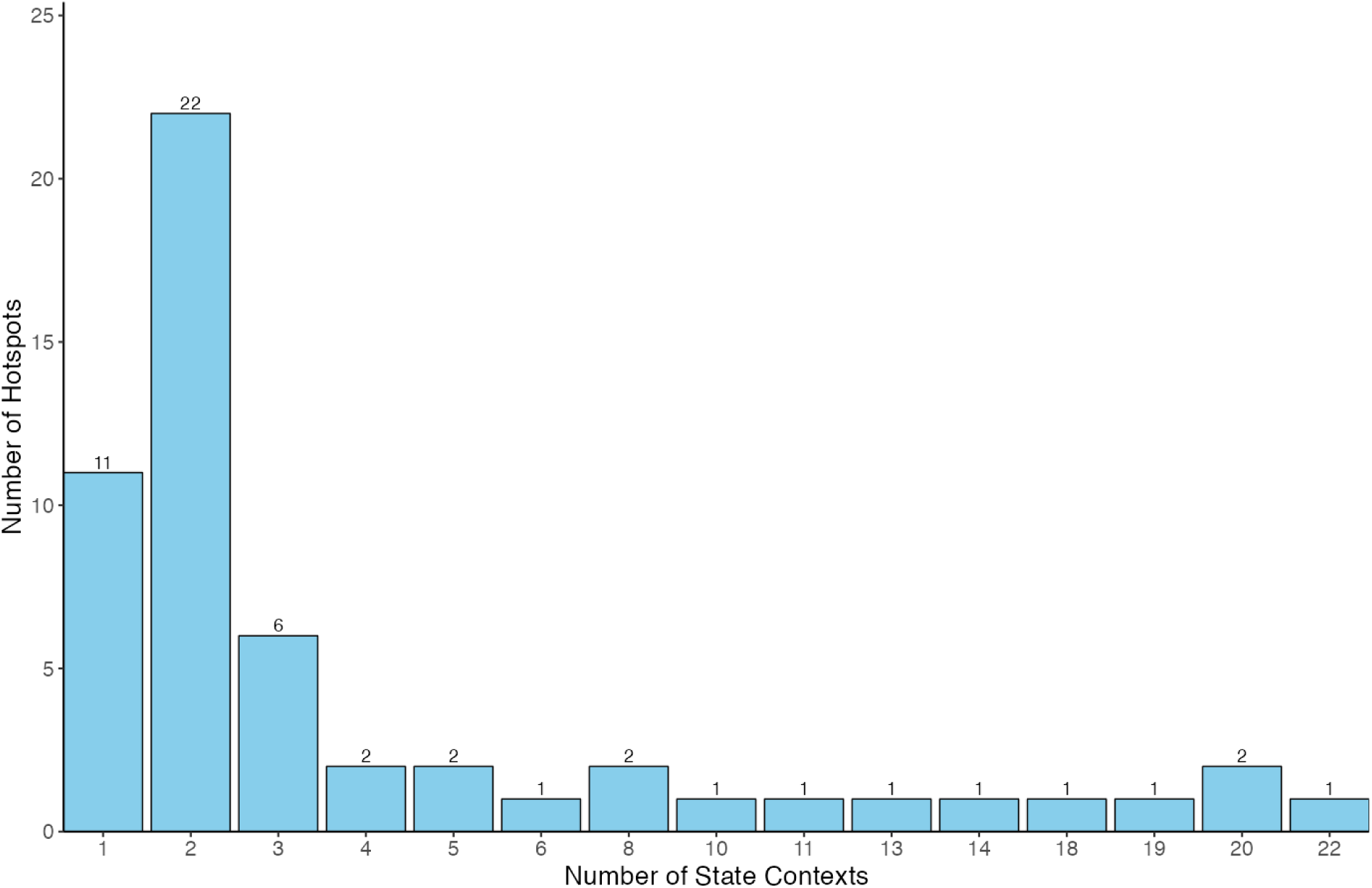
Histogram representing the number of states in which hotspots are observed. A majority of hotspots are identified in only one or two state contexts and very few are observed in five or more. In cases where no hotspots are shared across a specific number of state contexts, entries are omitted.

eQTL hotspots often overlap with fitness QTL, suggesting that they may arise from the same causal variants, and that the effects of the variants on fitness may be mediated via their effects on gene expression^3,7^. Sixteen of the 44 eQTL hotspots identified in this study overlap at least one previously-identified fitness QTL^10^. Some hotspots that influence the expression of hundreds to thousands of transcripts overlap many fitness QTL. Some hotspots that influence the expression of fewer genes also overlap fitness QTL and may play more context-specific roles in linking genetic effects on gene regulation and on fitness. **Below, we explore the likely biological effects of the hotspots we identified, connections between effects on expression and fitness, and candidate genes and variants underlying the hotspots**.

## eQTL hotspots in exponential phase cultures

Most prior eQTL studies in yeast have been conducted using cells grown to mid-log phase in rich media. We identified 19 hotspots in this condition in the two crosses (9 in BYxRM and 10 in CBSxYJM). We identified several hotspots previously found in bulk studies in these crosses under similar conditions, including those at *IRA2* and *MKT1* in the BYxRM cross and at *CYR1* in the CBSxYJM cross^8,11,14^. Consistent with prior work on interactions between eQTL hotspots and physiological states, we identified a hotspot on chromosome VIII that influenced expression of mating-related genes in the BYxRM cross in cells that occupied the G1 cell cycle phase^14^. This locus also influenced occupancy in the G1 phase state. One of these hotspots overlaps a fitness QTL identified in this cross.

## eQTL hotspots after acute salt perturbation

We identified 21 hotspots in samples collected 30 minutes after addition of salt (12 in BYxRM and 9 in CBSxYJM). Seven of these hotspots were also observed during exponential phase growth prior to the addition of salt. Six of these hotspots overlapped fitness QTL, three overlapped physiological state occupancy QTL, and four overlapped ESR gene set activity QTL (Table 3). These hotspots influence the expression of transcripts related to key components of the osmostress response including ribosomal gene regulation and protein folding, as well as sterol metabolism^32^.

**Table 3.** High-level summary of hotspots observed in the BYxRM salt experiment and overlaps with relevant QTL mapped in the same cross. Asterisks indicate that segregating variants with notable PROVEAN scores exist in the coding regions of indicated genes. Here, combined refers to hotspots identified at the level of cross:timepoint vs. cross:timepoint:state.

| Hotspot | BYxRM NaCl T0 | BYxRM NaCl T30 | Gene sets enriched | QTL Overlap | Hypotheses | Ontology |
| --- | --- | --- | --- | --- | --- | --- |
| chrI-I |  | G1 (27), G2_M_mating (56), Combined (132) | Glycolysis, peroxisome | ESR | OAF1*, GPB2* | TF. Cam/PKA regulator |
| chrII-I |  | Combined (41) | Translation, Trehalose anabolism |  | CYC8*, SUS1*, CMD1 | TF, TF, Calmodulin; Ca2+ binding protein |
| chrII-II | Combined (28) |  | Glutamate biosynthesis |  | IRA1* | GTPase-activating protein (gtpase regulator) |
| chrV-I | Combined (30), G2_M_mating (16) | Combined (87), G1 (94) | Ribosome | ESR | FAA2*, GPA2* | Medium chain fatty acyl-CoA synthetase, alpha subunit of the heterotrimeric G protein |
| chrV-II | S (16), combined (33) |  | Sulfate assimilation pathway II |  | CEM1 | beta-keto-acyl synthase |
| chrVII-I |  | Combined (66), G2_M_no_mating (33) | Translation | ESR, Fitness, State | GCN1*, MDS3 | Signaling pathway regulator, signaling pathway regulator (TOR) |
| chrVIII-I | Combined (35), G1 (22) |  | Conjugation with cellular fusion, pheromone response | Fitness, State | GPA1* | Subunit of G protein |
| chrXI-I |  | Combined (96) | Cytoplasmic translation |  | STE3* | Receptor for a factor pheromone |
| chrXII-I | Combined, many | Combined, many | Sterol biosynthesis, cellular respiration | Fitness | HAP1* | TF |
| chrXII-II |  | G1 (36) | Translation, Trehalose anabolism |  | CDC25* | guanine nucleotide exchange factor (gtpase regulator) |
| chrXIII-I | Combined, many | Combined, many | Oxoacid metabolic process | Fitness | PHO84*, GTR1 | Ion transporter, gtpase |
| chrXIII-II |  | Combined (105) | Purine metabolism |  | RPM2* | RNase subunit |
| chrXIV-I | Combined, many | Combined, many | Translation | Fitness | KRE33 | Ribosomal biogenesis protein |
| chrXIV-II | Many | Many | Translation, glycolysis | ESR, Fitness, State | MKT1* | Unclear/RNA binding protein |
| chrXV-I | Combined, many | Combined, many | Cellular respiration | Fitness, State | IRA2*, HAL9* | Signaling pathway regulator, TF |

### eQTL hotspots in stationary phase after 1 week

We identified seven hotspots in the BYxRM mapping panel that spent one week in stationary phase. Linking transcripts as well as putative causal variants were generally related to growth control and metabolic alternatives to glycolysis. One hotspot influenced glutamate biosynthesis genes (Table 4). Two hotspots influenced salvage pathway gene sets. One hotspot influenced transcripts enriched for sugar transporter gene sets, which could also be a response to starvation. One hotspot was enriched for oxidative phosphorylation gene sets and another was enriched for gene sets related to translation. Three of these hotspots overlapped fitness QTL, two overlapped a physiological state occupancy QTL, and four overlapped ESR gene set activity QTL (Table 4).

**Table 4.**
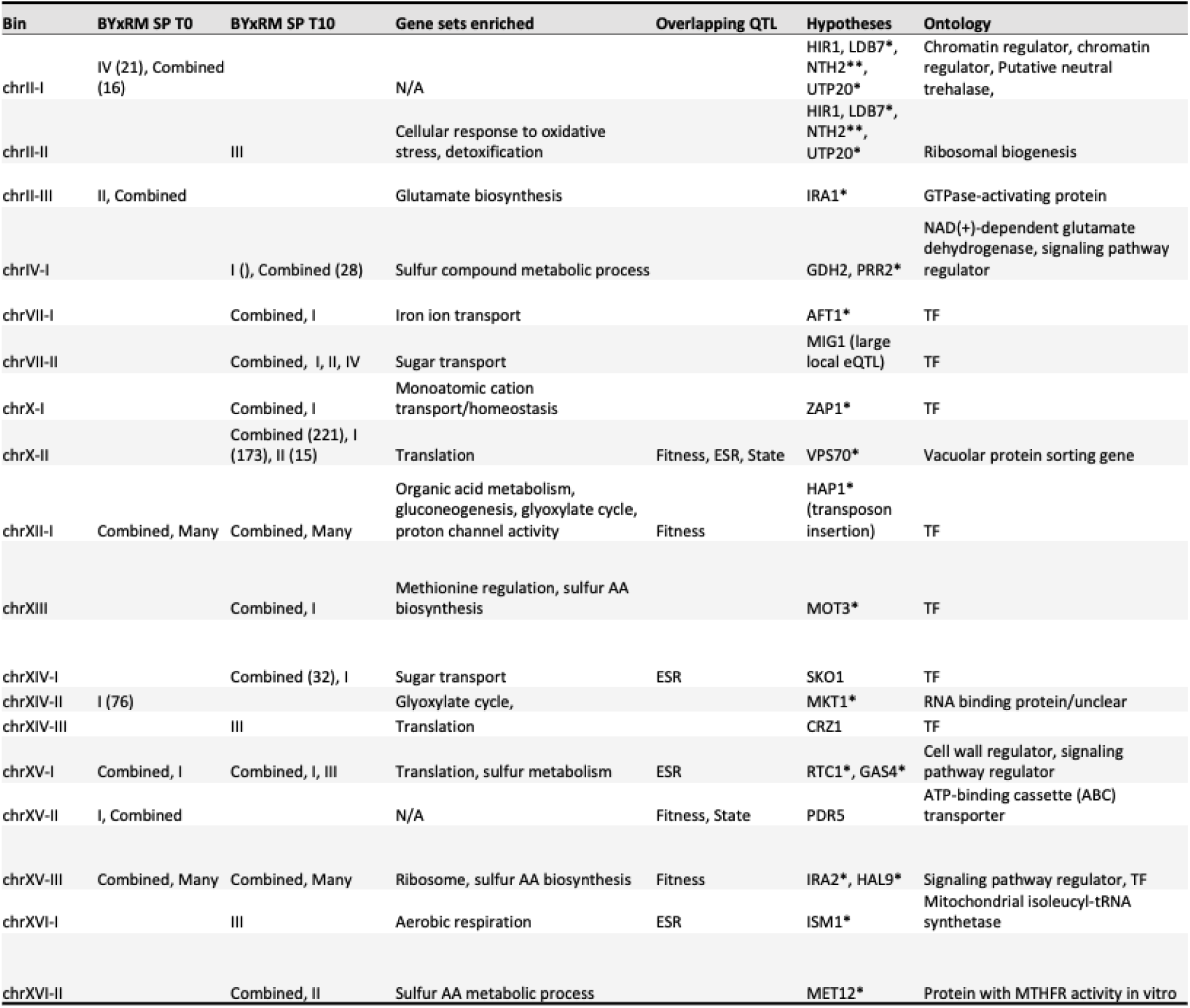
Hotspots observed in the stationary phase experiment and overlapping QTL. Asterisks indicate that segregating variants with notable PROVEAN scores exist in the coding regions of indicated genes.

| Bin | BYxRM SP T0 | BYxRM SP T10 | Gene sets enriched | Overlapping QTL | Hypotheses | Ontology |
| --- | --- | --- | --- | --- | --- | --- |
| chrII-I | IV (21), Combined (16) |  | N/A |  | HIR1, LDB7*, NTH2**, UTP20* | Chromatin regulator, chromatin regulator, Putative neutral trehalase, |
| chrII-II |  | III | Cellular response to oxidative stress, detoxification |  | HIR1, LDB7*, NTH2**, UTP20* | Ribosomal biogenesis |
| chrII-III | II, Combined |  | Glutamate biosynthesis |  | IRA1* | GTPase-activating protein |
| chrIV-I |  | I ( ), Combined (28) | Sulfur compound metabolic process |  | GDH2, PRR2* | NAD(+)-dependent glutamate dehydrogenase, signaling pathway regulator |
| chrVII-I |  | Combined, I | Iron ion transport |  | AFT1* | TF |
| chrVII-II |  | Combined, I, II, IV | Sugar transport |  | MIG1 (large local eQTL) | TF |
| chrX-I |  | Combined, I | Monoatomic cation transport/homeostasis |  | ZAP1* | TF |
| chrX-II |  | Combined (221), I (173), II (15) | Translation | Fitness, ESR, State | VPS70* | Vacuolar protein sorting gene |
| chrXII-I | Combined, Many | Combined, Many | Organic acid metabolism, gluconeogenesis, glyoxylate cycle, proton channel activity | Fitness | HAP1* (transposon insertion) | TF |
| chrXIII |  | Combined, I | Methionine regulation, sulfur AA biosynthesis |  | MOT3* | TF |
| chrXIV-I |  | Combined (32), I | Sugar transport | ESR | SKO1 | TF |
| chrXIV-II | I (76) |  | Glyoxylate cycle, |  | MKT1* | RNA binding protein/unclear |
| chrXIV-III |  | III | Translation |  | CRZ1 | TF |
| chrXV-I | Combined, I | Combined, I, III | Translation, sulfur metabolism | ESR | RTC1*, GAS4* | Cell wall regulator, signaling pathway regulator |
| chrXV-II | I, Combined |  | N/A | Fitness, State | PDR5 | ATP-binding cassette (ABC) transporter |
| chrXV-III | Combined, Many | Combined, Many | Ribosome, sulfur AA biosynthesis | Fitness | IRA2*, HAL9* | Signaling pathway regulator, TF |
| chrXVI-I |  | III | Aerobic respiration | ESR | ISM1* | Mitochondrial isoleucyl-tRNA synthetase |
| chrXVI-II |  | Combined, II | Sulfur AA metabolic process |  | MET12* | Protein with MTHFR activity in vitro |

## eQTL hotspots observed ten minutes after rich media perturbation

We identified 14 hotspots in the stationary phase perturbation time point. The hotspots reflect survival of starvation as well as processes necessary for transition back to the cell cycle and growth. Several of these hotspots influence transcripts related to nutrient transport (distinct hotspots influence import of sugars, sulfur, and iron) and utilization as well as regulation of metal ions. Two hotspots influenced stress response transcripts and two influenced translation-related transcripts. Four of these hotspots overlapped fitness QTL, one overlapped physiological state occupancy QTL, and four overlapped ESR gene set activity QTL (Table 4).

### Candidate causal genes and variants underlying the hotspots

Our examination of the hotspots uncovered a diverse set of candidate causal genes with a range of mechanistic hypotheses to explain their effects on gene expression (Tables 3 and 4). The most straightforward hypotheses involve coding variation or local eQTL effects that alter the function or expression of transcription factors, leading to downstream effects on the expression of their regulatory targets. Hotspots that fit this pattern include those we hypothesize are driven by variants in transcription factors *MOT3*, *AFT1*, and *CRZ1*, among others (Figure 11). Another class of hypotheses involves variants in genes that encode upstream regulators of signaling pathways. Candidate causal genes that could drive hotspots via this mechanism include GTPase regulators, GPCRs, and non-GPCR membrane-bound sensors, among others. Specific examples of candidate genes in this category include *GPA1*, which encodes a G protein subunit involved in pheromone response, as well as the HOG pathway osmosensor *HKR1*^14^. Other candidate genes belong to additional functional categories including chromatin regulators, ribosomal biogenesis factors, cell wall proteins, and transporters.

A majority of our hypotheses involve missense coding variation but there are examples of hotspots for which causal genes are likely influenced by local eQTL. One such hypothesis involves a local eQTL that influences the MIG1 transcription factor (TF) (LOD 72). The transcripts linking to this locus are enriched for gene sets related to sugar metabolism and transport. *MIG1* mediates glucose repression and is known to be environmentally responsive^30^.

. A majority of our causal hypotheses involve rare variants, but this study adds to a growing set of common variants that very likely drive expression hotspots and consequently fitness. One example is a common variant with a large predicted variant effect in *GPA1* that is causal for an eQTL hotspot as well as a cell cycle occupancy trait QTL^14,33^. We hypothesize that a common, missense variant in the zinc finger domain of the *MOT3* TF is causal for an eQTL hotspot observed in the CBSxYJM salt perturbation dataset as well as overlapping fitness and ESR gene set activity QTL (Figures S33, S34). In the case of the hotspot driven by *GPA1*, it is thought that the high allele frequency of the causal variation reflects balance of growth control and mating efficiency^14^. The high allele frequency of the hypothesized causal variant in *MOT3* discussed above could reflect a similar balance between induction of the environmental stress response and proliferation.

### Causal connections between hotspots, fitness QTL, and state QTL

We identified several hypotheses regarding causal relationships between expression traits and fitness. An NaCl t30 hotspot in CBSxYJM with linking transcripts enriched for sterol metabolism and sulfur metabolism gene sets (20 at the combined level) was identified on chrXIII. *MOT3*, a transcription factor situated at this locus, regulates sterol metabolism transcripts and is likely a causal gene. A common coding variant in *MOT3* (AF of 0.36) with a large computationally-predicted effect (PROVEAN score of –8.9) is likely the primary driver of the linkages to this hotspot. This variant occurs in a zinc finger domain. A fitness QTL for growth in YPD was fine-mapped to *MOT3* in this cross, further supporting the causal role of *MOT3* variation^10^.

The following example illustrates a causal hypothesis in which the mechanistic link between the candidate gene and the linking transcripts is less direct. A hotspot on chrX with linkages enriched for translation-related transcripts was observed in the SP t10 time point (Table 4). It overlaps several QTL for fitness in stressful conditions, one of which was fine-mapped to *VPS70* (‘vacuolar protein sorting 70’)^10^. We hypothesize that *VPS70* is also causal for the expression hotspot. This expression hotspot overlaps QTL for occupancy in states ‘I’ and ‘IV’, which are distinguished by higher expression of transcripts in the GO categories of “growth” and “translation-related transcription” and lower expression of stress-related genes. It is known that vacuolar function plays an important role in establishing, maintaining, and exiting quiescence, and it has been shown that deletion of *VPS70* increases chronological lifespan in the BY background^17,34,35^. A variant in *VPS70* that alters its role in vacuolar function could plausibly alter the expression of these gene sets and influence fitness, although the precise mechanism is currently unclear.

## Discussion

We conducted eQTL and physiological state QTL mapping in large yeast crosses subjected to acute perturbations using a single-cell RNAseq protocol. Our design involved acute salt perturbation of cells growing rapidly in rich media as well as acute nutrient repletion of one week old stationary phase cultures. eQTL mapping was performed with awareness of major state populations. Occupancy in specific states was also mapped in a separate analysis. The stationary phase and nutrient repletion time points included states that have not, to our knowledge, been characterized in depth or at all. We focused on eQTL hotspots, which are thought to explain a majority of expression trait heritability in all cases and are especially interpretable in the context of acute perturbation time points. We developed a host of hypotheses regarding causal relationships between the genetics of gene expression and fitness by overlaying expression and state QTL identified in this study with fitness QTL identified in the same crosses in prior work.

Expression QTL hotspots can influence the expression levels of sets of genes involved in the same pathways or cellular processes, thereby affecting cellular physiology and fitness. Prior work has shown the strong context-dependence of hotspot effects across states within an environment and our use of single-cell RNAseq here allowed us to observe context dependence across states within an environment and across environments^11^. Because of the one-pot nature of these experiments, we were able to incorporate acute perturbations in our design and accordingly mitigate some of the interpretive limitations of steady-state eQTL mapping. We observed that hotspot effects often depend on the physiological state in which they are mapped and can shift within 10-30 minutes after acute perturbations. This is much faster than demonstrated in prior GxE eQTL work, which was constrained by bulk methods and steady state measurement. The hotspots mapped after perturbations tended to influence transcripts involved with adaptation to new environmental conditions. Processes influenced by hotspots mapped after salt perturbation include protein folding, translation, and sterol metabolism, and examples drawn from the nutrient repletion experiment include transporter activity, oxidative stress, and translation.

A major goal of eQTL mapping is to develop causal hypotheses linking regulatory variation to other traits. In this study, we focused on connecting eQTL hotspots to state occupancy and fitness differences under stress conditions and during the sharp environmental transitions that yeast frequently encounter outside the laboratory. Toward this goal, we identified overlaps between eQTL hotspots, state QTL, and fitness QTL, many of which had been fine-mapped to single genes^10^. We prioritized candidate causal genes and variants using computational predictions of deleterious missense mutations and functional annotations of genes within QTL confidence intervals. One example drawn from the salt perturbation experiment is a missense variant in the zinc-finger domain of the *MOT3* transcription factor, that likely drives a hotspot enriched for sterol metabolism transcripts and consequently influences fitness in stress conditions. From the stationary phase perturbation experiment we identified a missense variant in *VPS70*, a less well-characterized gene, that may underlie a hotspot enriched for translation-related transcripts and affect fitness in stressful conditions.

Because we are using biparental yeast crosses, it is possible to study the effects of rare and common variants in the broader yeast population with the same statistical power. While many of our hypotheses involve rare, missense variants in coding regions driving hotspots, some implicate common variants. For example, the putative causal missense variant in the zinc-finger domain of *MOT3* is common in the yeast population and appears to drive a hotspot. Experimental validation of this hypothesis would add to a growing set of such observations^14^.

In a complementary analysis, we identified major physiological state populations in each sample and quantified continuous ESR gene set activity for individual cells. In the stationary phase samples, one or more of the identified populations likely correspond to quiescent cells. We conducted genetic mapping using state occupancy as well as the continuous ESR activity values as quantitative traits and observed that the resulting QTL frequently overlapped eQTL hotspots and fitness QTL. These overlaps suggest widespread causal relationships between regulatory variation, physiological state, and fitness^36^. They also illustrate the value of genetic mapping of gene set activity and similar traits, particularly in systems where distant eQTL are difficult to map using conventional methods^37,38^. Such approaches can effectively ‘short-list’ candidate loci that may function as eQTL hotspots prior to experimental validation.

Our one-pot design enabled high-resolution mapping of eQTL and state QTL across acute perturbation time courses in multiple crosses, generating a large set of hypotheses about the causal basis of regulatory and fitness QTL. Although this study was powered to detect eQTL hotspots and small-to-moderate-effect distant eQTL, an increase in the numbers of cells per time point from thousands to millions would improve power and resolution. Because we destructively sampled cells, we could not definitively assign functional identities to all state populations. In the case of cellular quiescence, a particularly important state and one that is arguably easiest to study in yeast, future studies can build on our results, sorting and functionally characterizing stationary phase subpopulations, and directly testing their abilities to return to the cell cycle after nutrient repletion. Our results highlight the importance of eQTL hotspots in understanding the mechanistic basis of genetic effects on fitness and argue for continued attention to distant eQTL and state QTL, as well as for the development of methods to carry out similar approaches in multicellular systems.

## Materials and Methods

### Sampling cultures for single-cell RNAseq

At appropriate points in each experiment, aliquots were taken from culture flasks and concentrated on nylon filters via a vacuum apparatus. Filters were placed into 50mL conical tubes which were submerged in a mixture of dry ice and ethanol. Tubes were subsequently transferred to –80C for storage prior to processing.

### Segregant single-cell RNAseq experiments

Diploids were streaked out, grown, and put through the random spore prep as in Boocock at al (2025)^14^. Spores were treated with zymolyase 2000 U/mL (Amsbio, #120491-1) and inoculated in YPD. At mid-log, after just enough time for the markers to be expressed, segregants were separated according to mating type using FACS (Biorad s3e). We note that the sorting strategy is imperfect and a small fraction of MATalpha cells were carried into the rest of the experiment. MATa cells were grown overnight and allowed to reach stationary phase after the 1mL or so of sorter fluid with 250k segregants post-sorting was diluted with fresh YPD. After growing overnight, segregants were spun down, media was replaced with new YPD, and at mid-log the experiments began with differences in the details of the perturbation. For the NaCl perturbation time courses, crystalline NaCl was added into the culture flasks to bring them to 0.7M NaCl as the means of perturbation. We did this to avoid conflating the effects of spinning down the culture with the specific response to changes in salt concentration. Another sample was taken thirty minutes after the perturbation. For the stationary phase experiment, segregants were processed the same way but after reaching mid-log, they were allowed to continue on to stationary phase and were kept in the shaker at 30C for 1 week. Sampling occurred after one week. Next, 1mL of the stationary phase culture was added to fresh YPD already at 30C to serve as the beginning of the nutrient repletion time course. Another sample was taken ten minutes later.

### Processing frozen samples prior to single-cell RNAseq library construction

Suspensions of cells at 1M cells/mL were prepared and provided to the UCLA TCGB core facility for 10x chromium 3’GEX library construction. In short, filters were removed from –80C and fixed with 80% methanol before being washed, sonicated, and diluted. We counted cells using a Bulldog Bio hemocytometer. Where appropriate, we sonicated samples that showed obvious clumping for periods of 1min. We did the minimal amount of sonication (duration and intensity) necessary to separate most cells based on visual inspection. The standard 10x Genomics 3’ V3 single-cell RNAseq library construction protocol (using our provided concentration of cells as an input and a target of 10k cells as the output) was followed with a single deviation in which zymolyase 2000 U/ml is added to the 10x reagent (in particular the beads)^39,40^.

### Batching

In all cases, timepoints from the same experiments were removed from –80C and processed together through library construction. Libraries corresponding to timepoints from the same experiment were sequenced on the same flowcells. Illumina Novaseq was used for all experiments.

### Processing of sequence data

We generated fastq files using bcl2fastq via the Illumina basespace portal. Cellranger was used to process raw fastq files and output an expression matrix as described in Boocock et al. (2025)^14^. Filtered expression matrices were read into Seurat and processed similarly to what is described in Boocock et al. (2025)^14,23,41,42^. In short, count distributions were visualized and filtered based on unexpected outliers (total UMI count for technical issues and mating gene expression to confirm a lack of MATalpha contamination in mapping panels). Transcripts were filtered if they were not measured in at least 5 cells in the sample they were derived from.

### Clustering analyses to assign discrete physiological state labels to single-cells

Sctransform was used for variance stabilization and PCA was done with either the entire filtered expression matrix or a subset of transcripts based on literature (Spellman et al. for mid-log cell cycle, for example)^23,43^. Leiden clustering was conducted using the top 12 PCs. Clusters were evaluated using marker gene expression (as much as possible, we used genes for which mRNA expression is itself interpretable) from the literature as well as cluster-specific marker genes identified by Seurat’s findmarkers tool^41,42,44^. With the goal of assigning a relatively small number of state labels to these sometimes continuous datasets, we collapsed similar clusters within experiments and attempted to apply labels with similar meaning across experiments (see SI for an extended discussion of this process). We note that some cell cycle markers originally identified using yeast cultures cycling in rich media or after alpha factor synchronization may not have the same interpretive value in our perturbed conditions. We also note that while the amount of MAT alpha contamination is low, the mating response may be on in some of the MATa cells that we use for these analyses as a consequence of pheromone-related signaling vs. the particular perturbations used in this study.

### Genotype inference in mapping panels

We used deep sequencing data for the parent strains and a hidden markov model to infer segregant genotypes. This was the same approach described in Boocock et al. (2025)^14^.

### Mapping state occupancy traits

In short, we considered occupancy in or outside of a particular physiological state assignment (cluster) as a binary trait and conducted genetic mapping using a logistic regression model. See Boocock et al. (2025) for additional details regarding physiological state (clustering-based) QTL mapping^14^.

### Assignment of continuous ESR gene set activity scores

We assigned ESR activity scores using the AUCell tool^45^. Briefly, the transcriptome of each single cell is ranked and enrichment scores are generated for query gene sets using the most highly-expressed genes in each cell. We used the ESR gene (RP, RiBi, iESR) sets described in Gasch et al. (2017)^22^. For this phase of the analysis, Monocle3 was used to process Cellranger output matrices prior to using AUCell but only the resulting gene set activity values were used in downstream analyses^46^.

### Mapping of ESR activity traits

ESR activity scores for each gene set within each cross by timepoint combination were used for genetic mapping. Correlation coefficients were calculated using the cross product of the design matrix and ESR activity phenotype data. These coefficients were scaled and transformed into LOD scores.

### eQTL mapping and expression QTL hotspot assignment

Individual expression traits were mapped in local and genome-wide scans using negative binomial regression. Mapping was conducted individually for each physiological state/clustering-based subset of cells within each time point. See Boocock et al. (2025) for full details regarding eQTL mapping^14^. Expression hotspots were assigned as in Boocock et al. (2025)^14^. Briefly, the genome was divided into 50kb bins and, per chromosome, the mean number of linkages per bin was used as lambda in a poisson model.

### Counting overping QTL

We visually inspected pileups of eQTL confidence intervals at hotspot loci and at points manually combined 50kb bins (see SI). We overlaid ESR and state QTL confidence intervals to manually count overlaps of QTL confidence intervals. In cases where ESR QTL confidence intervals were larger than 15kb, we truncated them to 7.5kb on either side of the central marker.

### Gene set enrichment testing

We used gprofiler2 in R to conduct all gene set enrichment testing^47^. In all cases, we use relevant custom backgrounds of transcripts that are smaller than the full yeast genome. We do this to mitigate overinflating the significance of our results. An example of a custom background is the set of filtered transcripts (5000 of 6000+ yeast transcripts, for example) that are used for a differential expression analysis.

### Homology Modeling

We used an existing homology modeling result available from InterPro for the MOT3 transcription factor to assess potential influences of a putative causal mutation (Pro377His)^48^.

## Supporting information

Supplementary Tables (Hotspot Enrichment)

## Acknowledgements

We thank Stefan Zdraljevic and Giancarlo Bruni for helpful manuscript feedback and edits. We thank the Kruglyak lab for helpful discussion throughout the project.

## Author contributions

NA, JB, and HM performed experiments with assistance from JSB. NA analyzed data with assistance from JSB. LK and JSB supervised the project. NA, JSB, and LK wrote the manuscript. All authors discussed and agreed on the final version of the manuscript.

## Data availability

eQTL mapping outputs and fine-mapping results are available on Zenodo at https://doi.org/10.5281/zenodo.19929439. Sequencing data, gene expression matrices, and links to additional post-processed materials will be shared in a revised version of this preprint.

## Code availability

Code for the HMM and eQTL mapping can be found at https://github.com/joshsbloom/single_cell_eQTL/tree/master/yeast_GxE/code. Code to recreate the figures in the manuscript can be found at https://github.com/noahalexander/eqtl.gxe.2026. This repository also contains the code that performs trans-eQTL hotspot analysis, state characterization, and raw data processing.

