## Supplementary Information for "The genetics of transcriptional responses to stress in yeast"

I. State QTL LOD plots

**Stationary phase BYxRM**

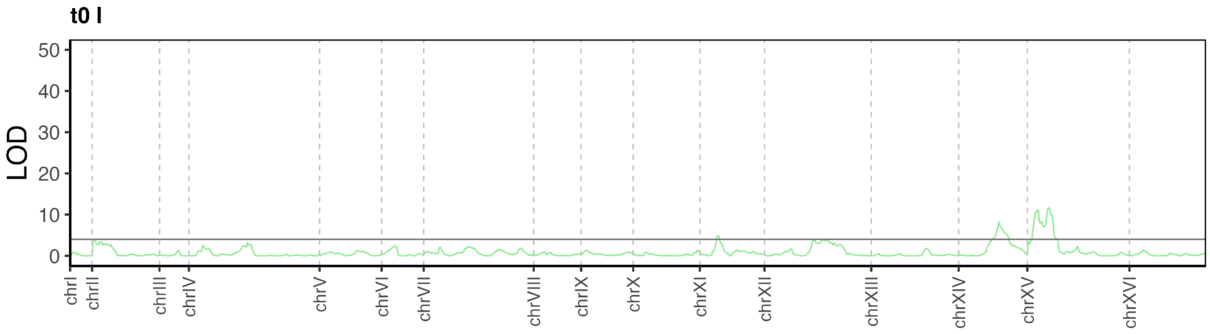


**Figure S1**. LOD trace of physiological state I occupancy mapping for the t0 timepoint of the BYxRM stationary phase experiment.

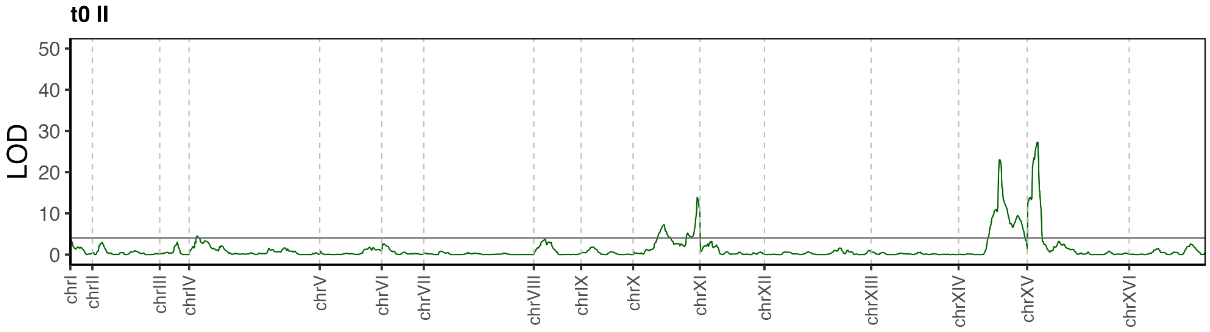


**Figure S2**. LOD trace of physiological state II occupancy mapping for the t0 timepoint of the BYxRM stationary phase experiment.

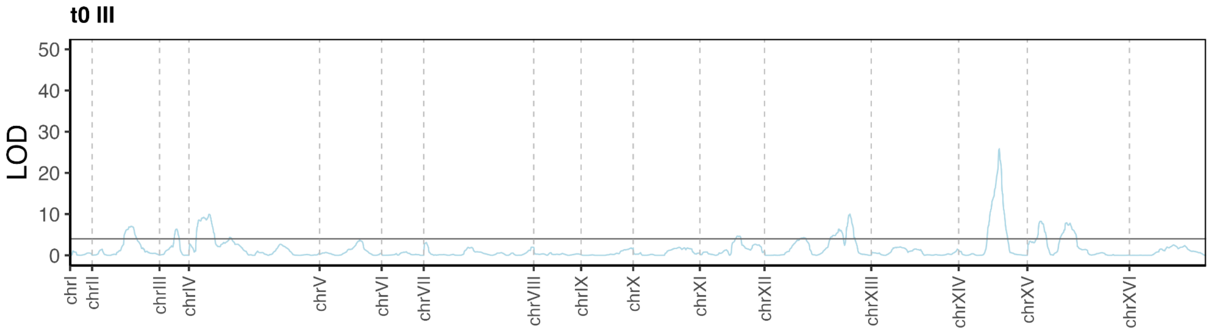


**Figure S3**. LOD trace of physiological state III occupancy mapping for the t0 timepoint of the BYxRM stationary phase experiment.

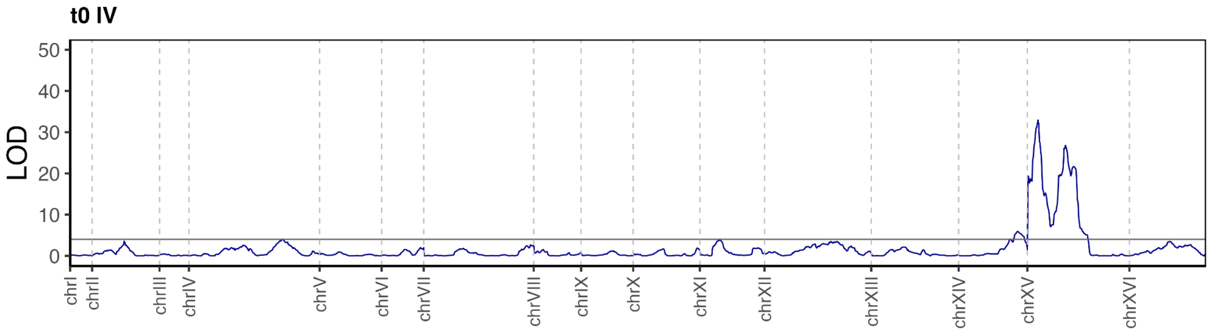


**Figure S4**. LOD trace of physiological state IV occupancy mapping for the t0 timepoint of the BYxRM stationary phase experiment.

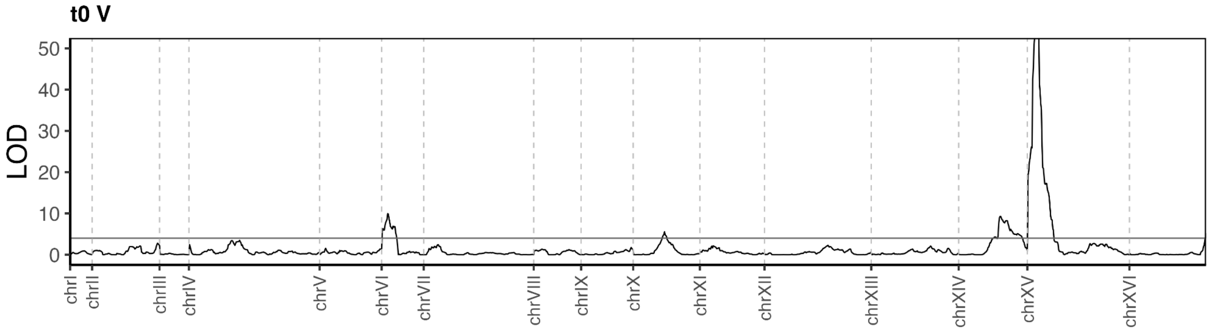


**Figure S5**. LOD trace of physiological state V occupancy mapping for the t0 timepoint of the BYxRM stationary phase experiment.

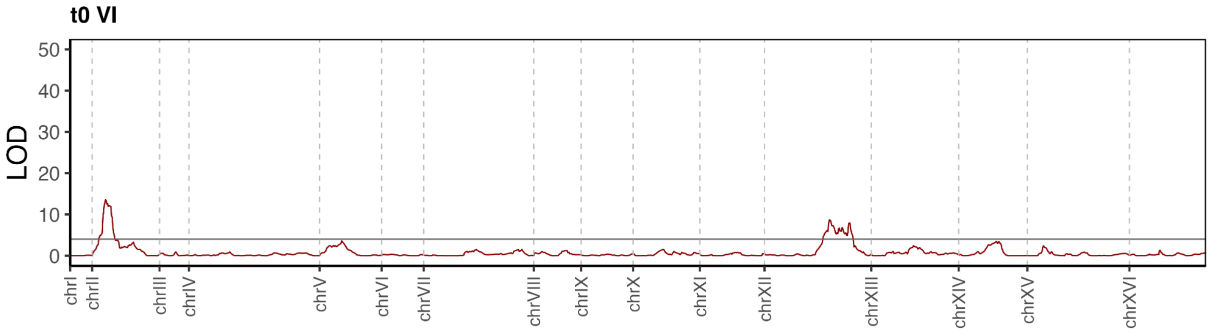


**Figure S6**. LOD trace of physiological state VI occupancy mapping for the t0 timepoint of the BYxRM stationary phase experiment.

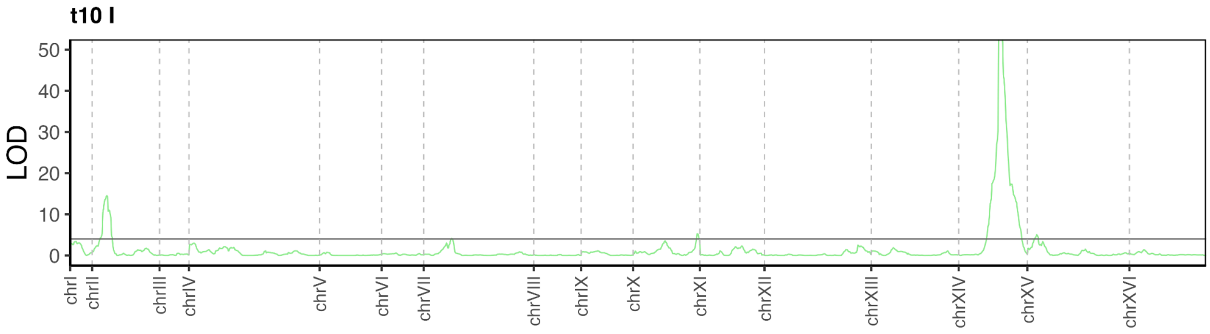


**Figure S7**. LOD trace of physiological state I occupancy mapping for the t10 timepoint of the BYxRM stationary phase experiment.

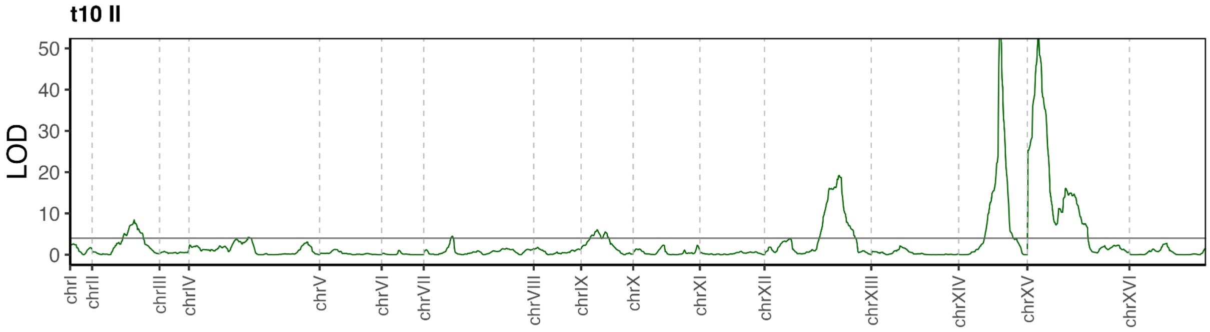


**Figure S8**. LOD trace of physiological state II occupancy mapping for the t10 timepoint of the BYxRM stationary phase experiment.

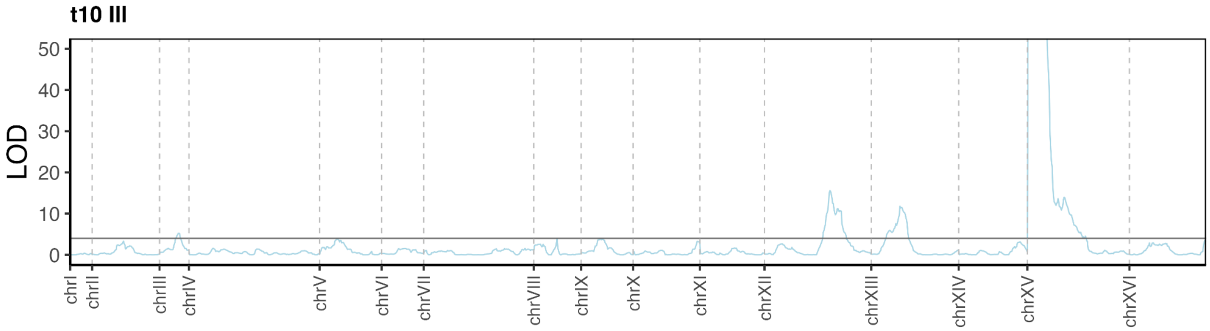


**Figure S9**. LOD trace of physiological state III occupancy mapping for the t10 timepoint of the BYxRM stationary phase experiment.

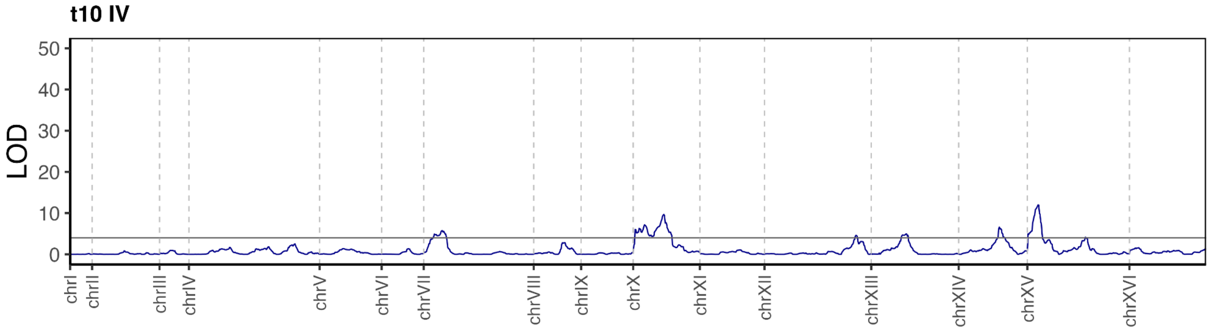

**Figure S10**. LOD trace of physiological state IV occupancy mapping for the t10 timepoint of the BYxRM stationary phase experiment.

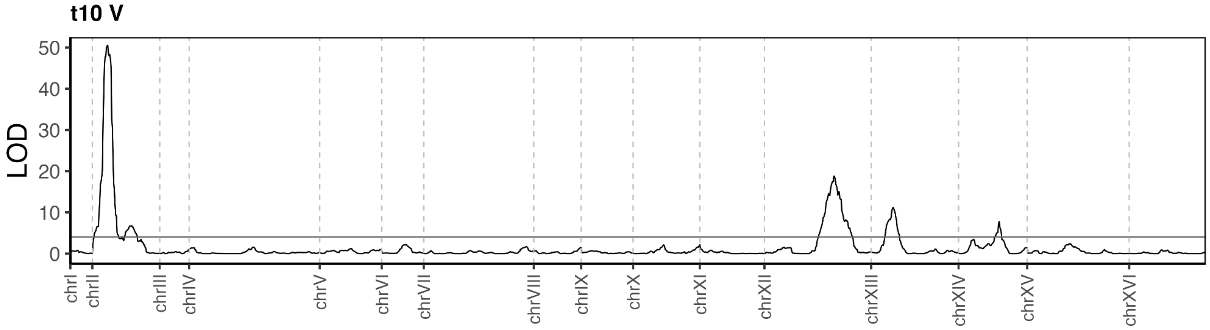


**Figure S11**. LOD trace of physiological state V occupancy mapping for the t10 timepoint of the BYxRM stationary phase experiment.

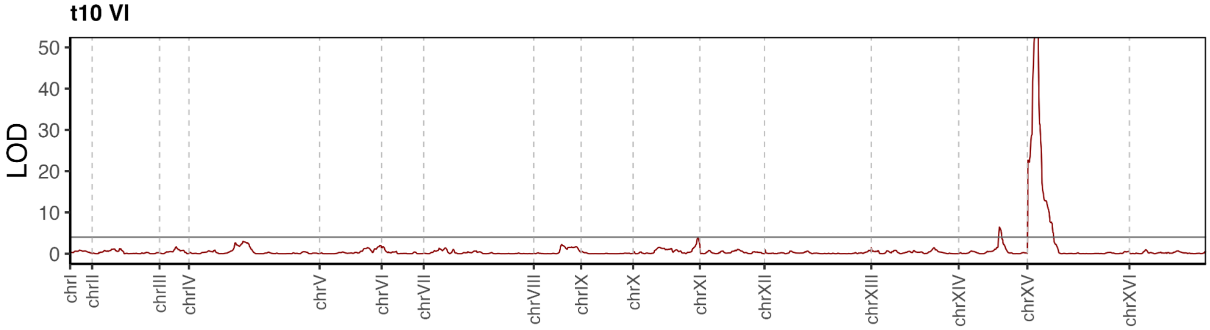


**Figure S12**. LOD trace of physiological state VI occupancy mapping for the t10 timepoint of the BYxRM stationary phase experiment.

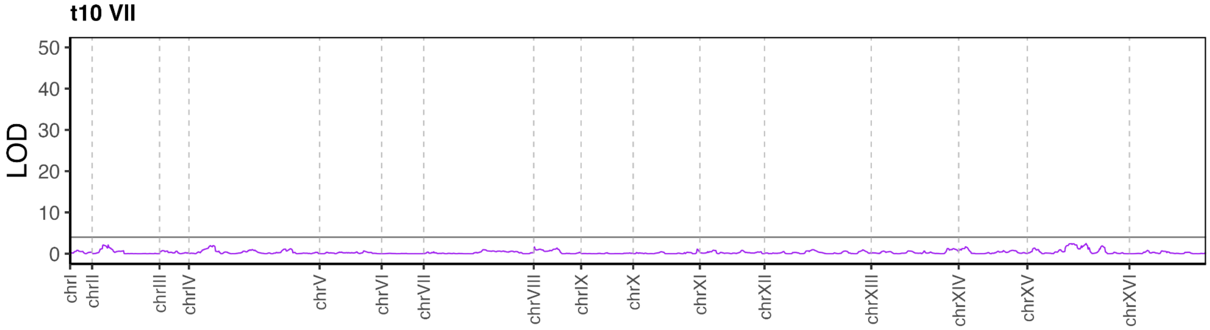


**Figure S13**. LOD trace of physiological state VII occupancy mapping for the t10 timepoint of the BYxRM stationary phase experiment.

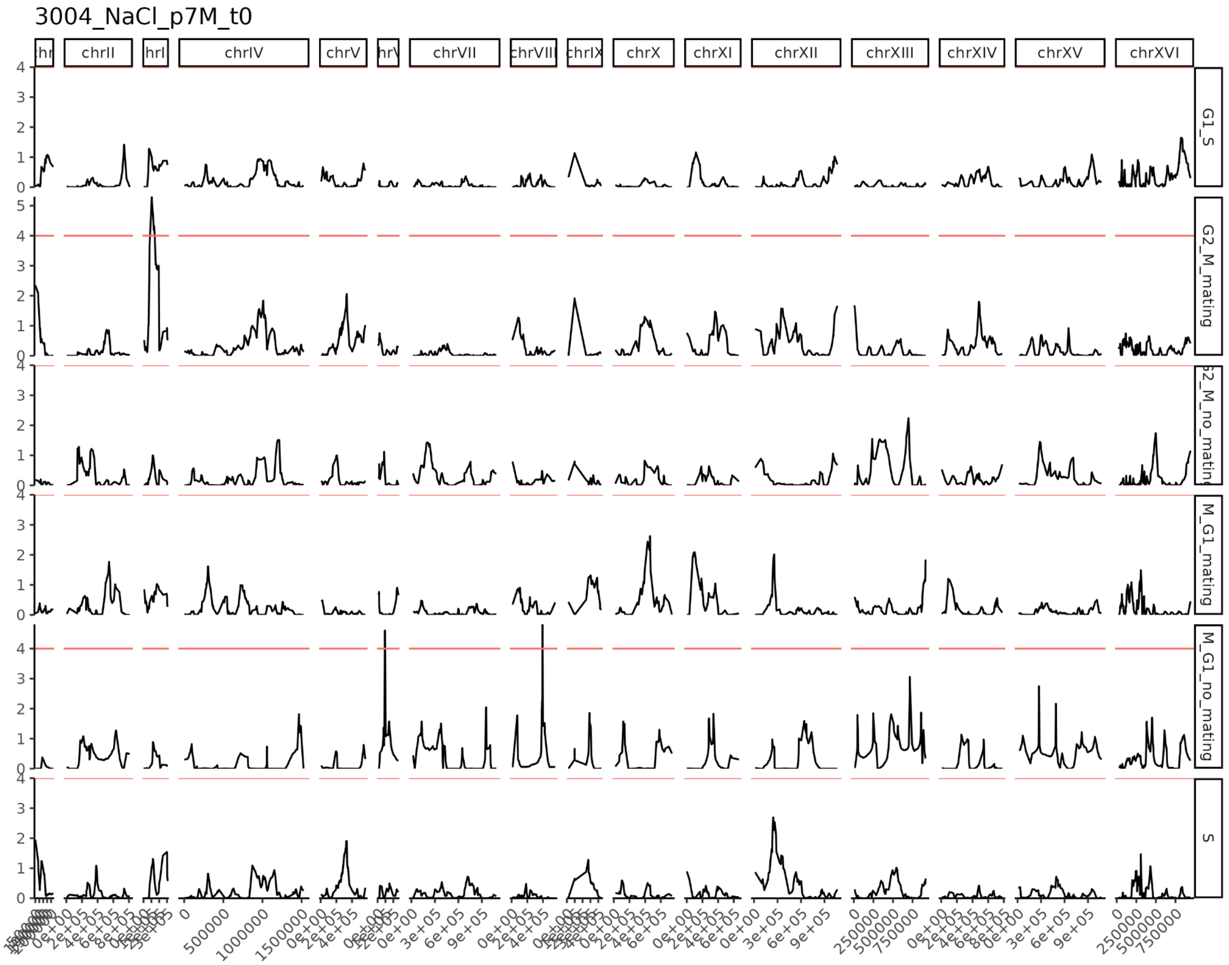


**Figure S14**. LOD trace of physiological state occupancy mapping for the t0 timepoint of the CBSxYJM salt perturbation experiment. Each row reflects a LOD trace reflecting mapping of occupancy in the state specified on the right.

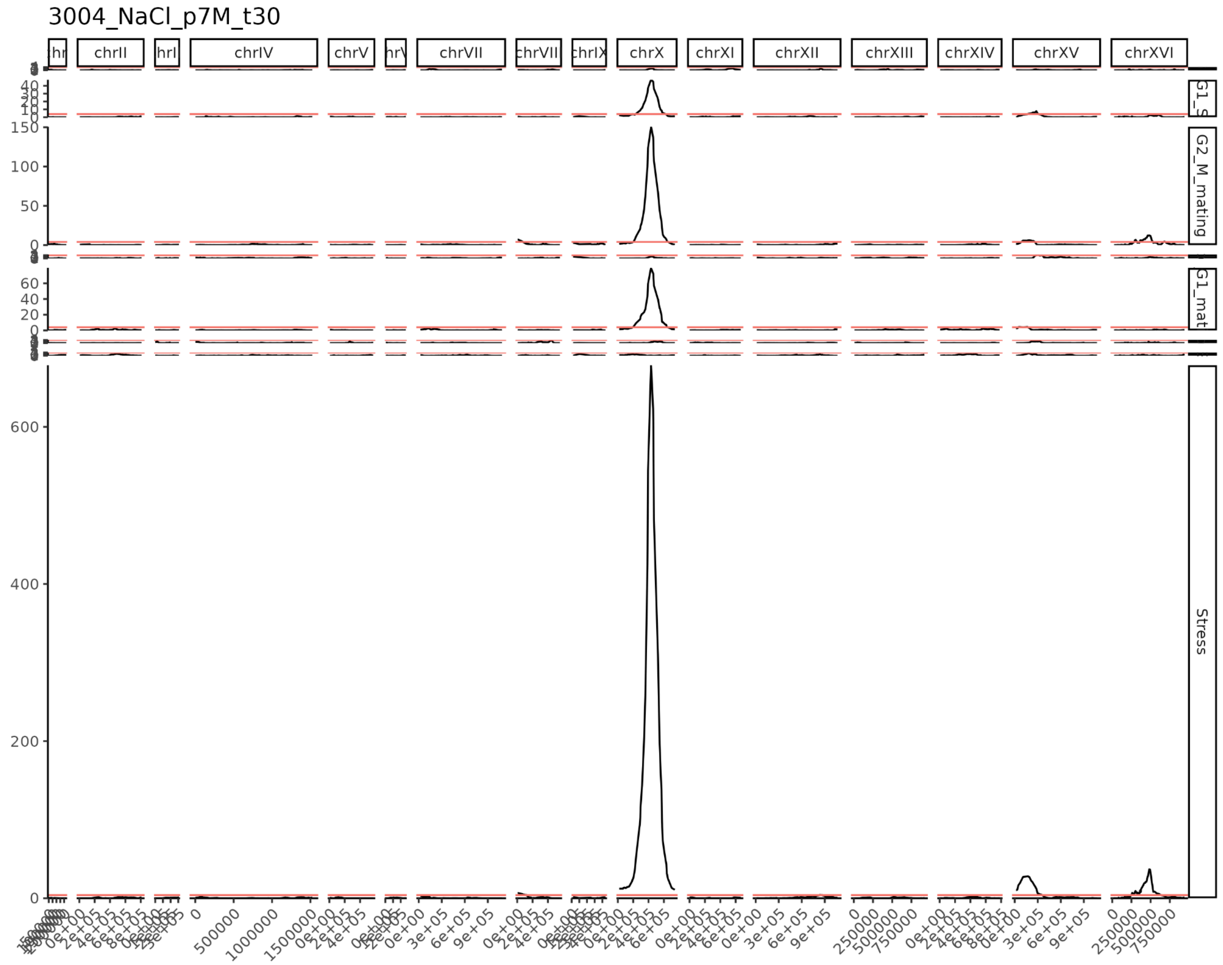


**Figure S15**. LOD trace of physiological state occupancy mapping for the t30 timepoint of the CBSxYJM salt perturbation experiment. Each row reflects a LOD trace reflecting mapping of occupancy in the state specified on the right.

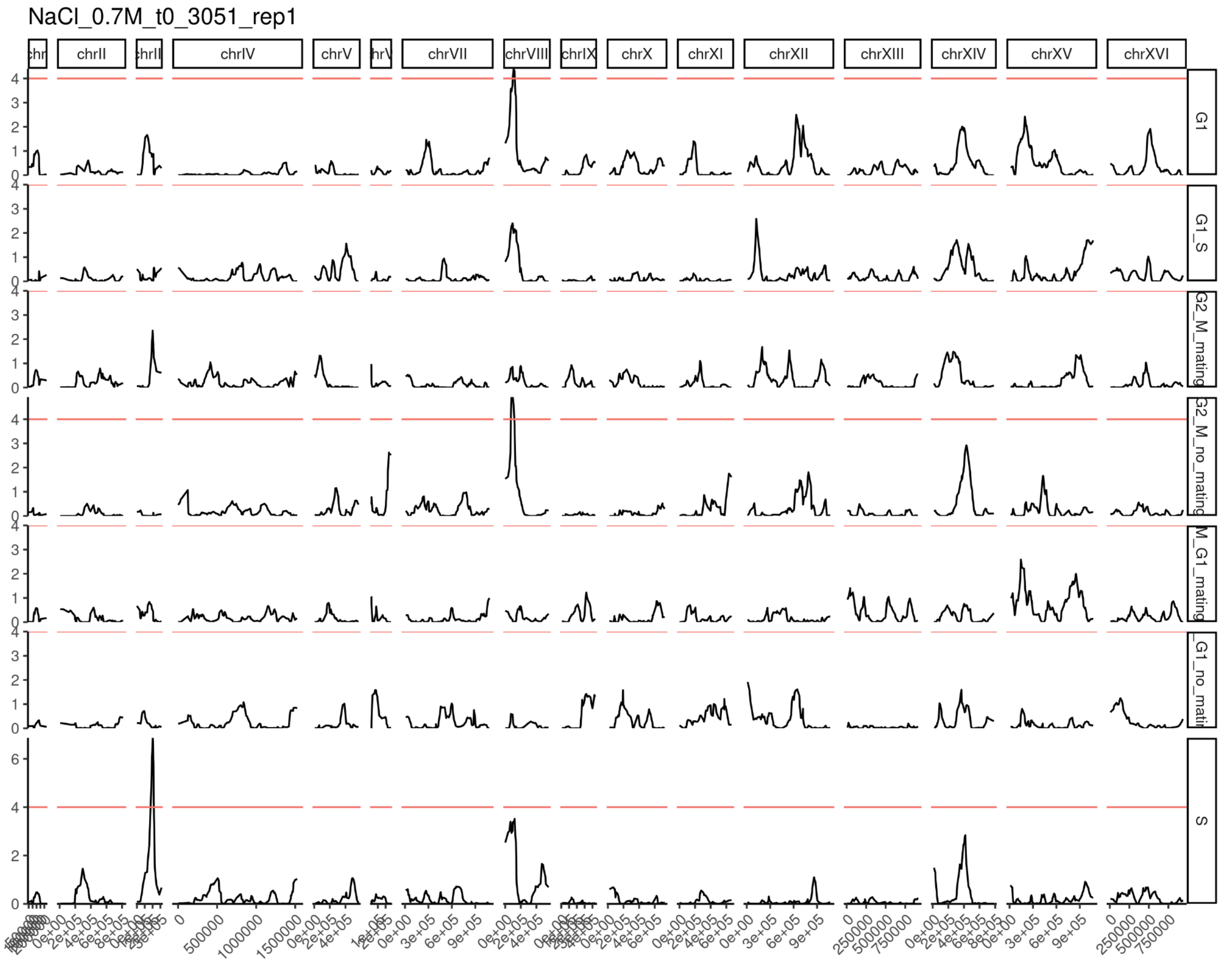


**Figure S16**. LOD trace of physiological state occupancy mapping for the t0 timepoint of the BYxRM salt perturbation experiment. Each row reflects a LOD trace reflecting mapping of occupancy in the state specified on the right.

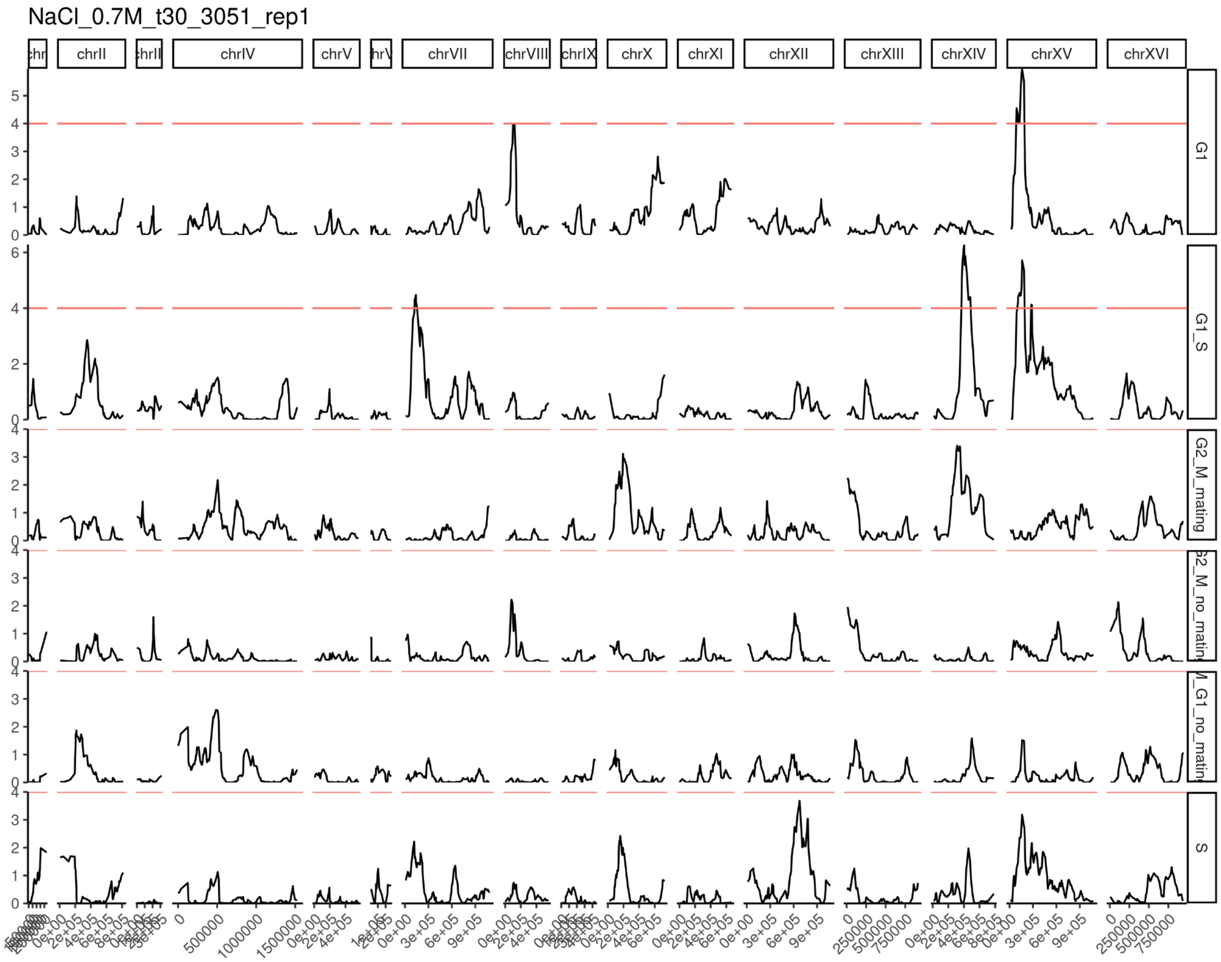


**Figure S17**. LOD trace of physiological state occupancy mapping for the t30 timepoint of the BYxRM salt perturbation experiment. Each row reflects a LOD trace reflecting mapping of occupancy in the state specified on the right.

II. ESR activity QTL LOD plots

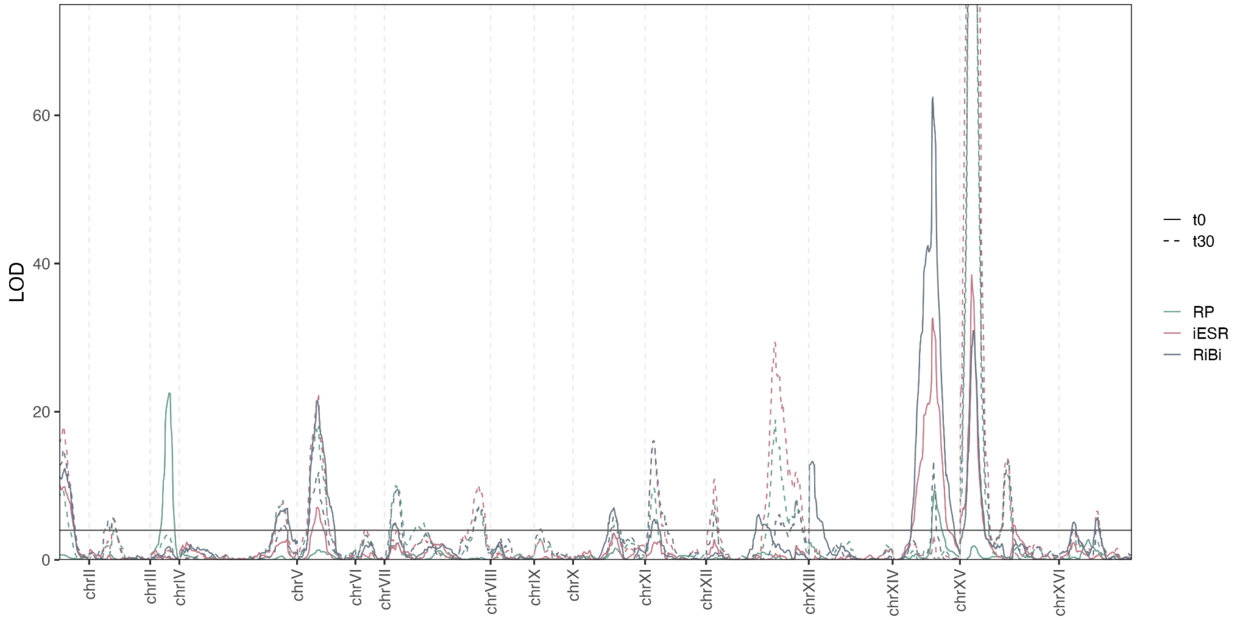


**Figure S18.** LOD traces for ESR activity traits in the BYxRM salt perturbation experiment. The colors reflect mapping results for distinct ESR gene set activity phenotypes. The continuous traces reflect results from the t0 time point and the dashed traces reflects mapping results for the t30 time point.

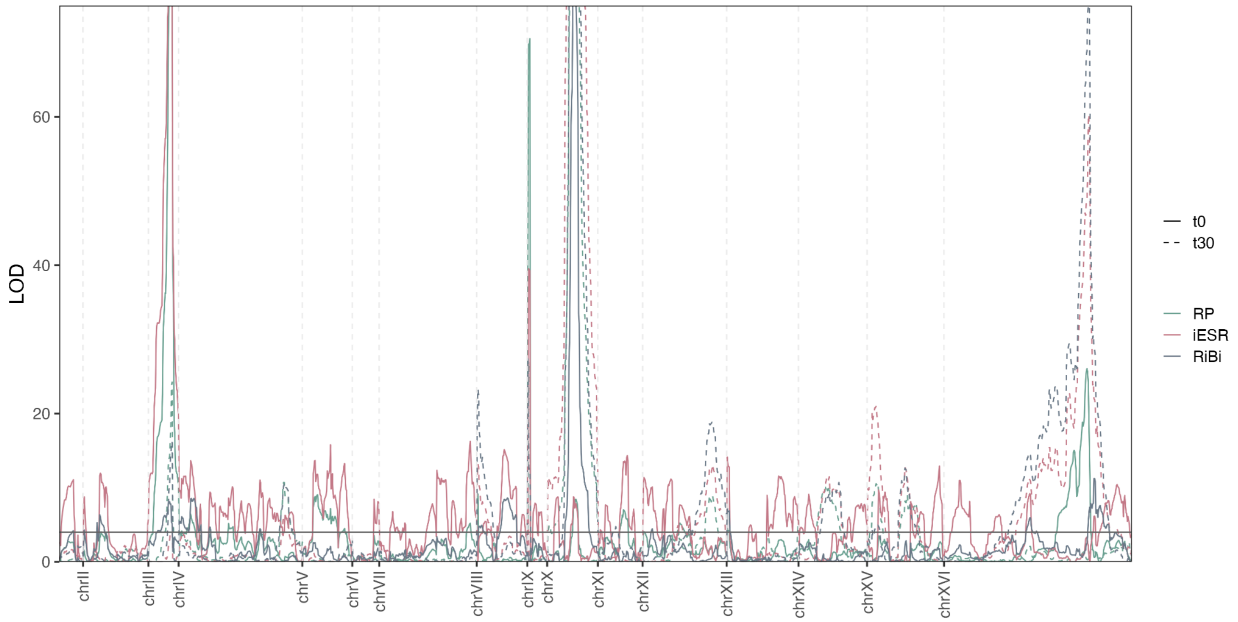


**Figure S19.** LOD traces for ESR activity traits in the CBSxYJM salt perturbation experiment. The colors reflect mapping results for distinct ESR gene set activity phenotypes. The continuous traces reflect results from the t0 time point and the dashed traces reflects mapping results for the t30 time point.

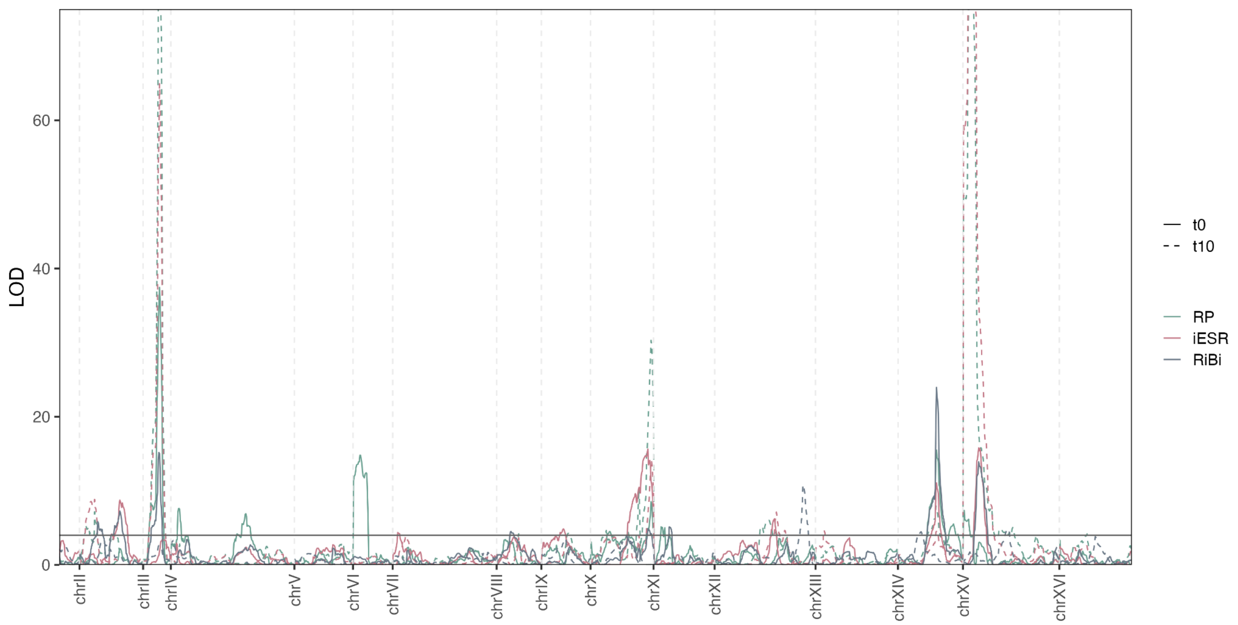


**Figure S20.** LOD traces for ESR activity traits in the BYxRM stationary phase experiment. The colors reflect mapping results for distinct ESR gene set activity phenotypes. The continuous traces reflect results from the t0 time point and the dashed traces reflects mapping results for the t30 time point.

III. Cis eQTL plots (t0 vs perturbation) for all experiments

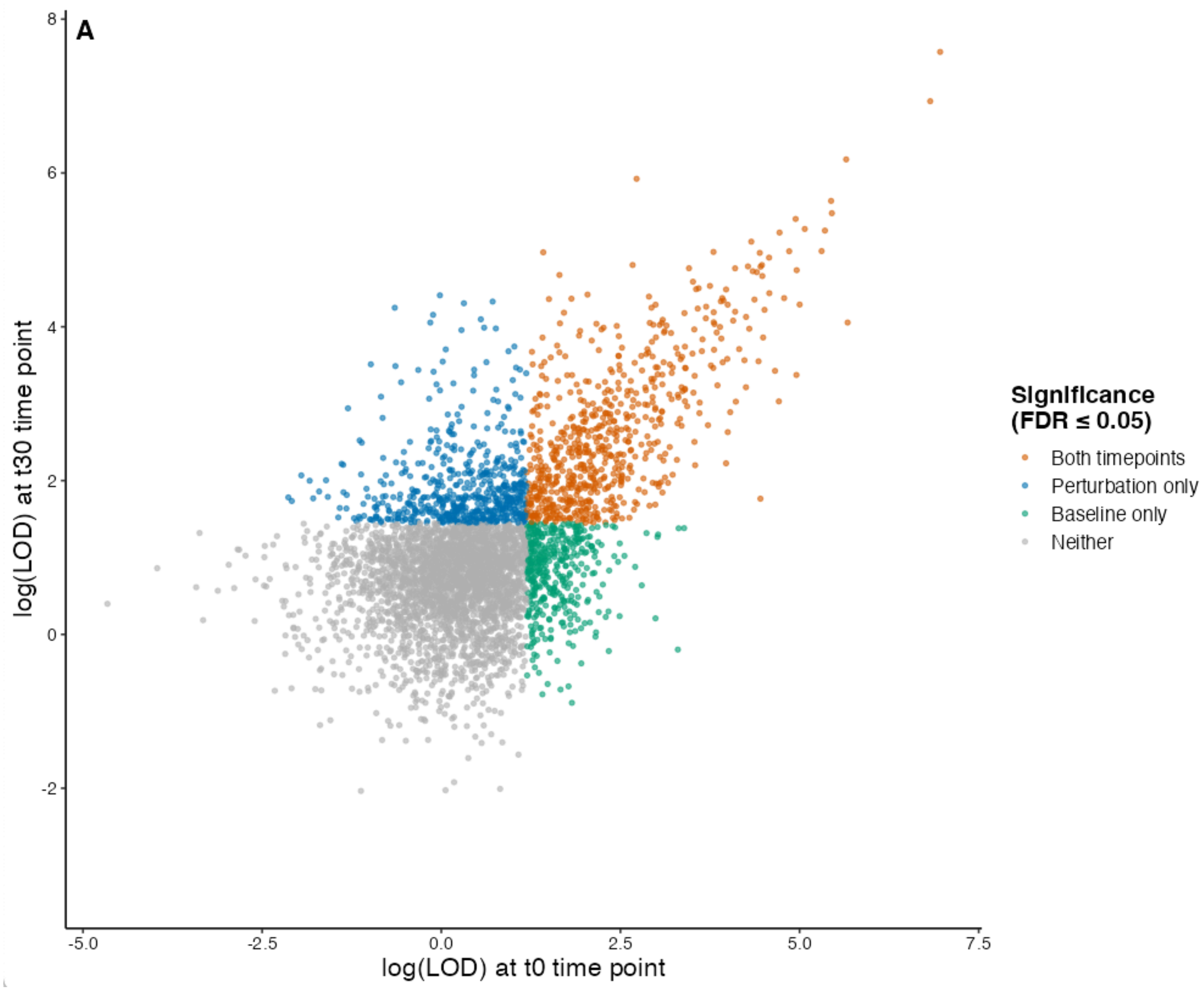


**Figure S21.** Local eQTLs mapped in the CBSxYJM salt perturbation experiment. The axes reflect LOD scores from the NaCl t0 and NaCl t30 conditions, respectively.

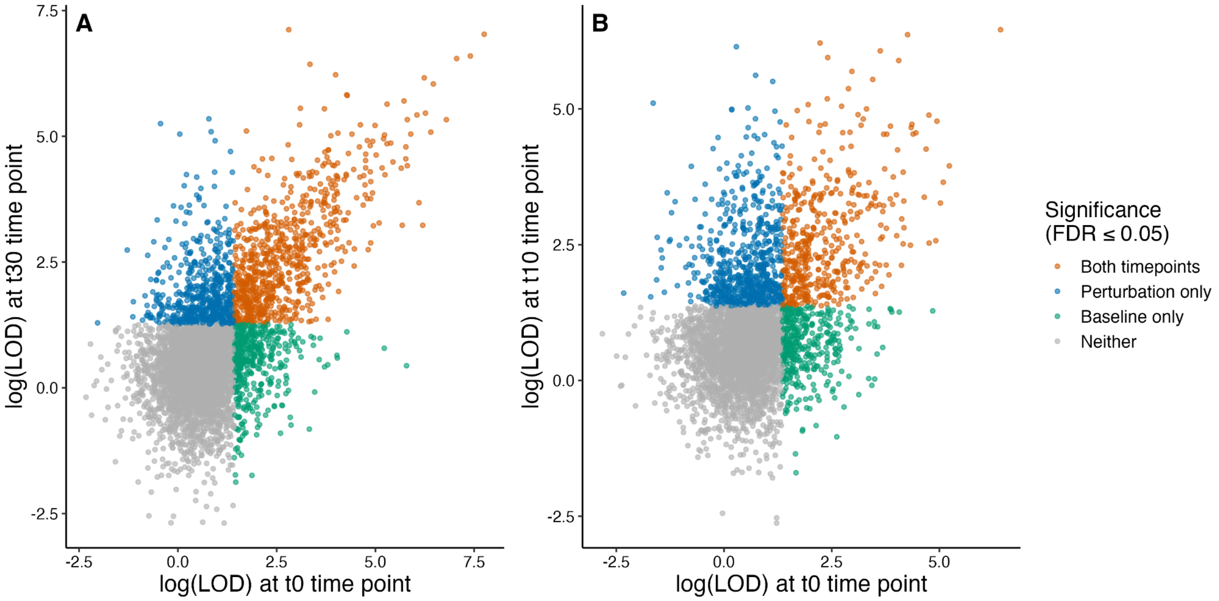


**Figure S22.** Local eQTLs mapped in the BYxRM perturbation experiment. Panel A shows LOD scores for transcripts corresponding to the NaCl t0 and NaCl t30 conditions. Panel B shows LOD scores for transcripts corresponding to the SP t0 and SP t10 conditions.

### IV. In-depth description of clustering-based state assignment in the salt experiments (stationary phase work is addressed in the body of the results section)

Our goals were to conduct clustering in PC space at high-resolution to characterize the heterogeneity in our samples and to assign a minimal set of state labels to each sample. We combined clusters where possible. We did this individually in each sample instead of analyzing every time point and/or cross together. We tried to use similar criteria for assigning the same labels across datasets. If we did not see evidence of a specific state we did not include it. We build upon the work of Spellman et al., but add a G1/S phase. We often use CLN2 as a G1/S marker in spite of it being labeled as a G1 gene in Spellman et al. Our CC markers were chosen to be as biologically interpretable as possible at the level of mRNA expression. These are often the genes with the strongest periodicity scores in Spellman et al.

#### BYxRM NaCl t0

Clusters 0 and 5 exhibited striking G2/M expression and were collapsed to a single g2/m phase. We note that cluster 0 has STE2 expression while cluster 5 does not. Regarding S phase, cluster 1 appears to share expression markers between G2/M and S but has much fainter expression of the band of S markers than cluster 3

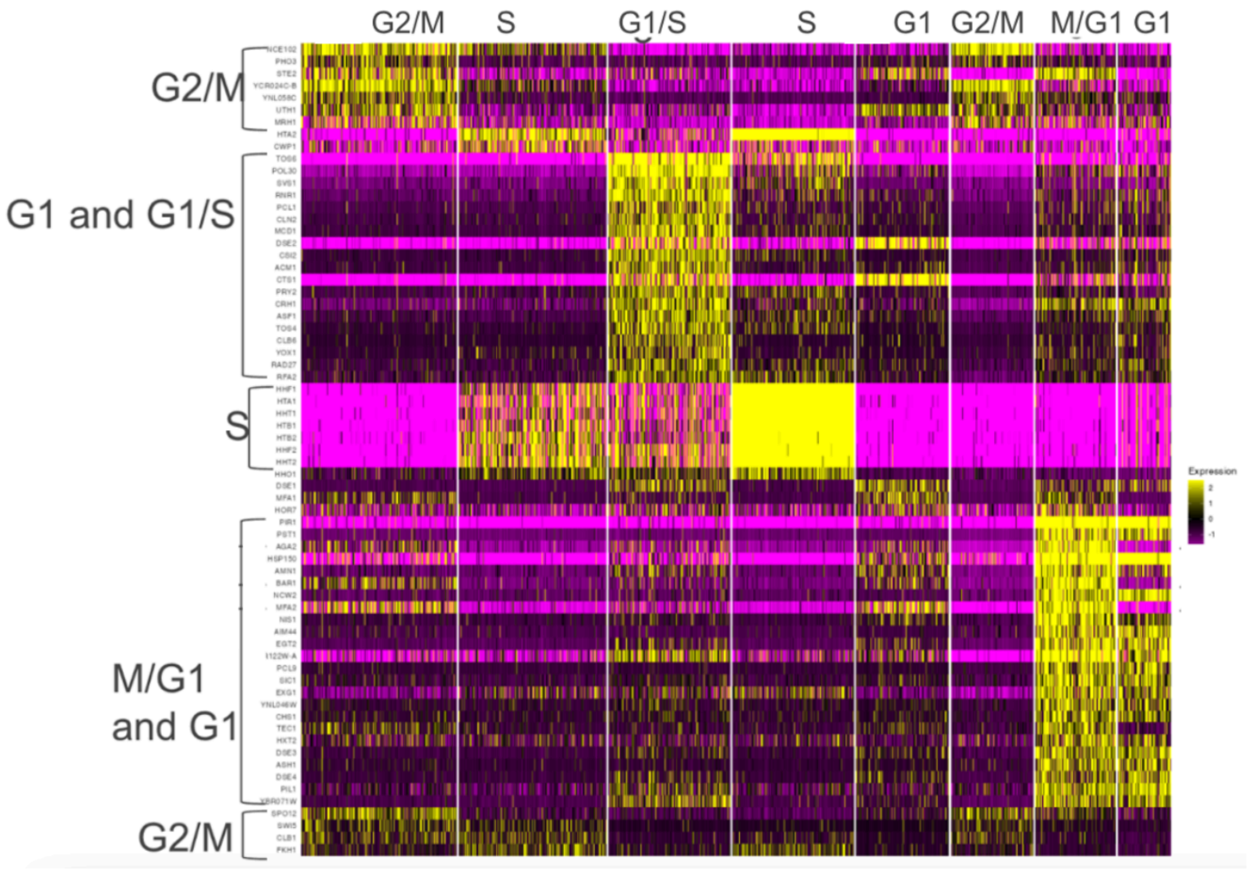


**Figure S23.** PCA-based clustering at high resolution in the BYxRM mid-log sample. G1 marker CTS1 can be seen most distinctly in cluster 4 while clusters 6 and 7 display characteristic M/G1 gene expression (PIR1, for example). Clusters 6 and 7 are distinguished by mating gene expression and are classified as m/g1 with mating genes and m/g1 without mating genes, respectively. As CTS1 and canonical G1 markers can be seen in 4 more distinctly than in any other cluster, we assign it G1. We note that in CBSxYJM NaCl t0, no cluster at this resolution has such distinct expression of CTS1.

Cluster 4 has much more muted expression of M/G1 genes and stronger expression of g1 marker cts1 than cluster 6. Focusing on M/G1 marker EGT2, for example, it is on in several cluster 4 cells but not nearly as strongly as in clusters 6 and 7. These clusters seem to both represent cells that are very early cell cycle yet are distinct with respect to mating gene expression (MFA2, for example). We have assigned labels ‘M/G1 + mating genes’ and ‘M/G1 without mating genes’ to 6 and 7 respectively.

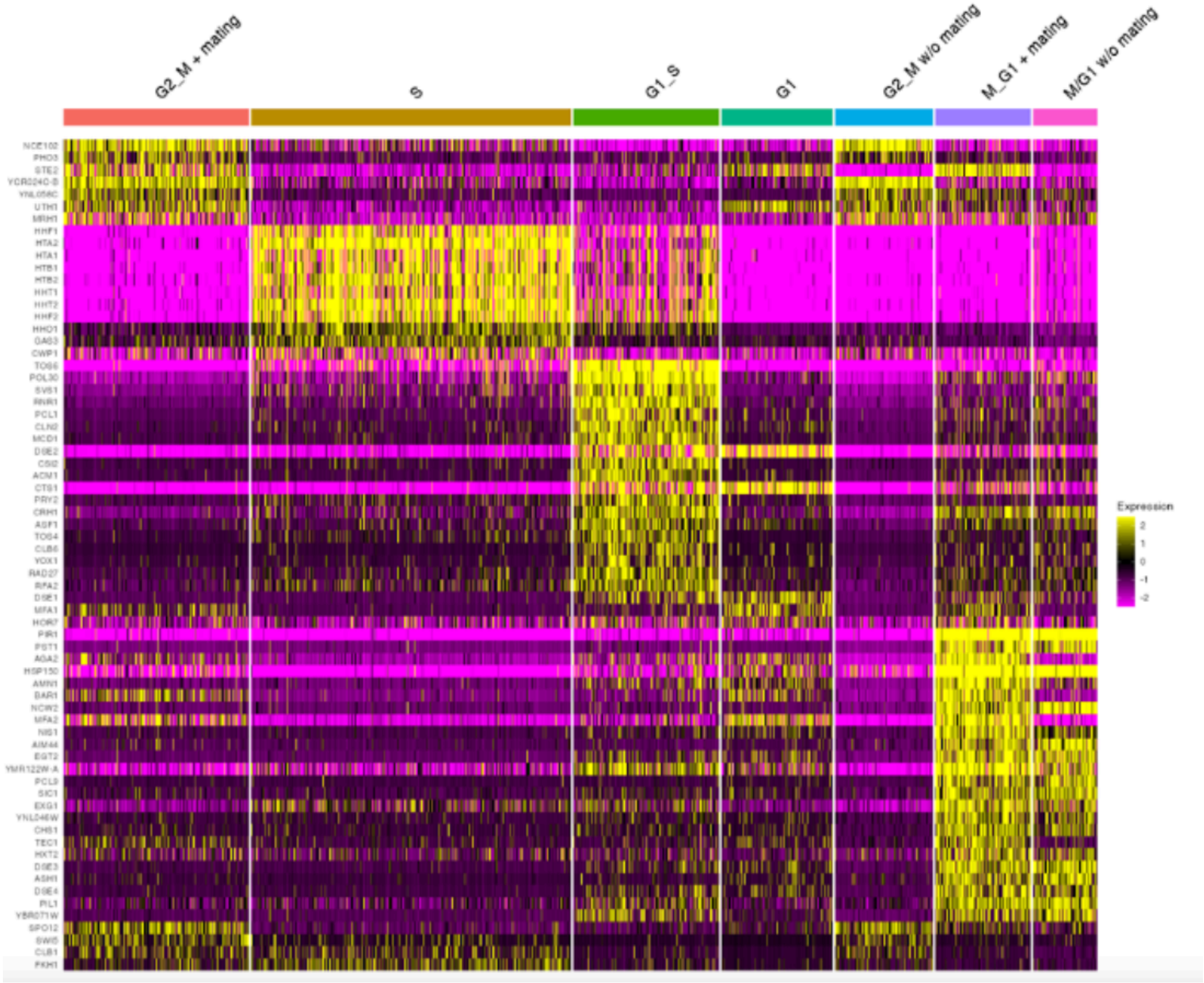


**Figure S24:** Collapsed state labels for BYxRM NaCl t0. Note the g1 cluster here does not have an analog (at least not one found at the higher end of clustering resolution we used/not with enough cells to conduct mapping) in the 3005 NaCl t0 dataset.

**BYxRM NaCl t30**

As with the t0 conditions, we tried several clustering resolutions and used one that generated enough clusters to capture the expected CC phases but was not so high that it generated clusters that were too small to use for mapping.

At high resolution, there are some small clusters with very extreme character that tend not to persist when clustering at lower resolution. In particular, the ‘stress’ cluster and ‘ribosomal protein’ cluster (at clustering res 0.55, see below) exhibit striking stress response related gene expression signatures, many related to osmotic stress. At 0.55 res clustering, many clusters have some degree of ‘stress’ character but it is diffuse/heterogeneous within each cluster except for cluster 8, which has very high expression of several stress-related transcripts.

Because the RP cluster/cluster 9 also had CC expression (S phase), we added this cluster to S but note that it is very distinct. Because the ‘stress’ cluster/cluster 8 did not exhibit much cc-related transcription at all relative to any other cluster, we left it as a distinct cluster. We also collapsed 9 with S since the number of cells in this cluster was very small (cluster 8 is not much smaller but we will keep it intact).

Clusters 1 and 7 are probably both g1 but at different parts of it or with different things still expressed (POL30, SVS1, for example). See CTS1 expression in both clusters.

Clusters 0 and 5 can be collapsed to G2/M but it seems like they have expression signatures from different parts of the g2/m transition.

Cluster 6 is somewhat distinct from the other G2/M clusters but we are deciding to compress it with the other putative G2/M.

Based on CLN2 expression and lower expression of canonical S markers, we assign cluster 2 as G1/S (a few other clusters have CLN2 expressed in most cells but it is nearly entirely off in many). This transcript is on in more cells than in mid-log YPD/NaCl t0 cultures.

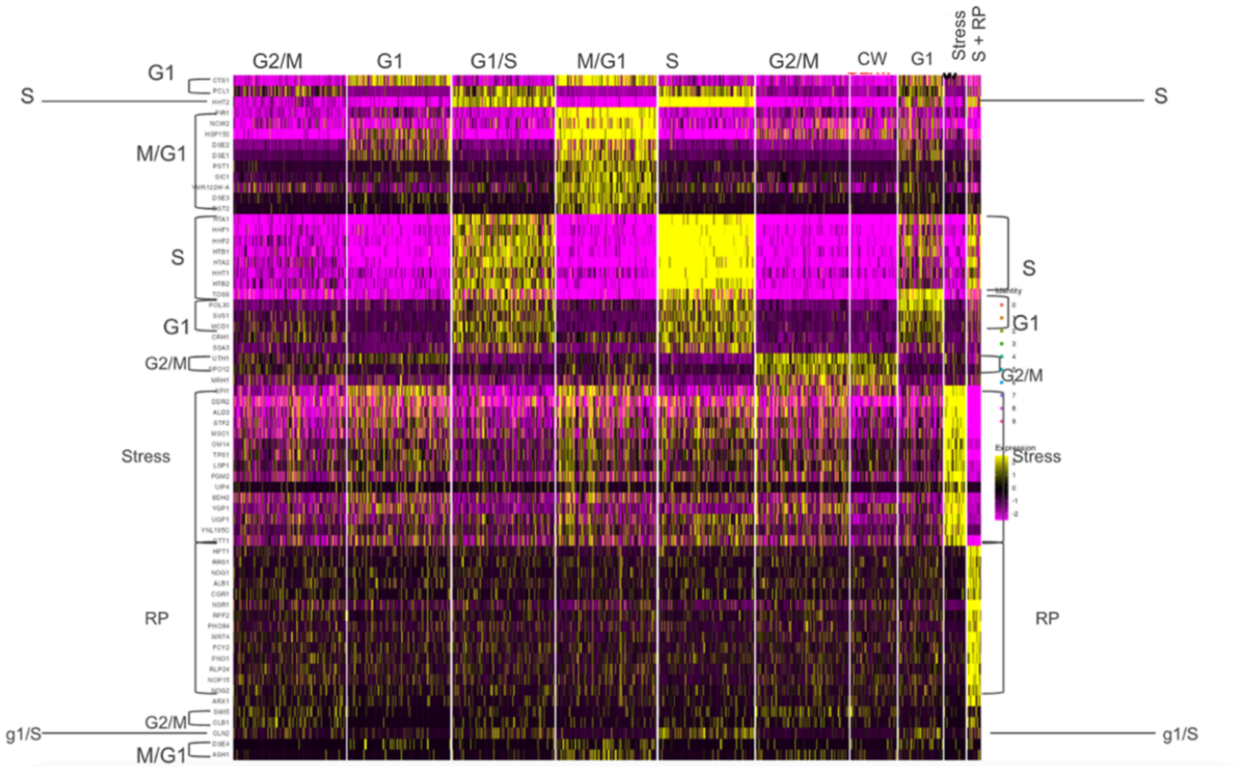


**Figure S25.** Clusters identified in 12 dimensional PC space separate the columns (no clustering of cells is done for this heatmap but the clusters are themselves separated). Z-scores are visualized for genes that are known CC markers as well as top marker genes for each cluster.

We do not have power to do mapping in sets of cells as small as the ‘stress’ cluster from 0.55 resolution and were not sure how to best place these cells within the conventional cell cycle. We thought that lower resolution clustering would be most appropriate and noticed that at resolution of 0.43, the stress cluster is no longer produced but there is sufficient nuance to assign CC labels.

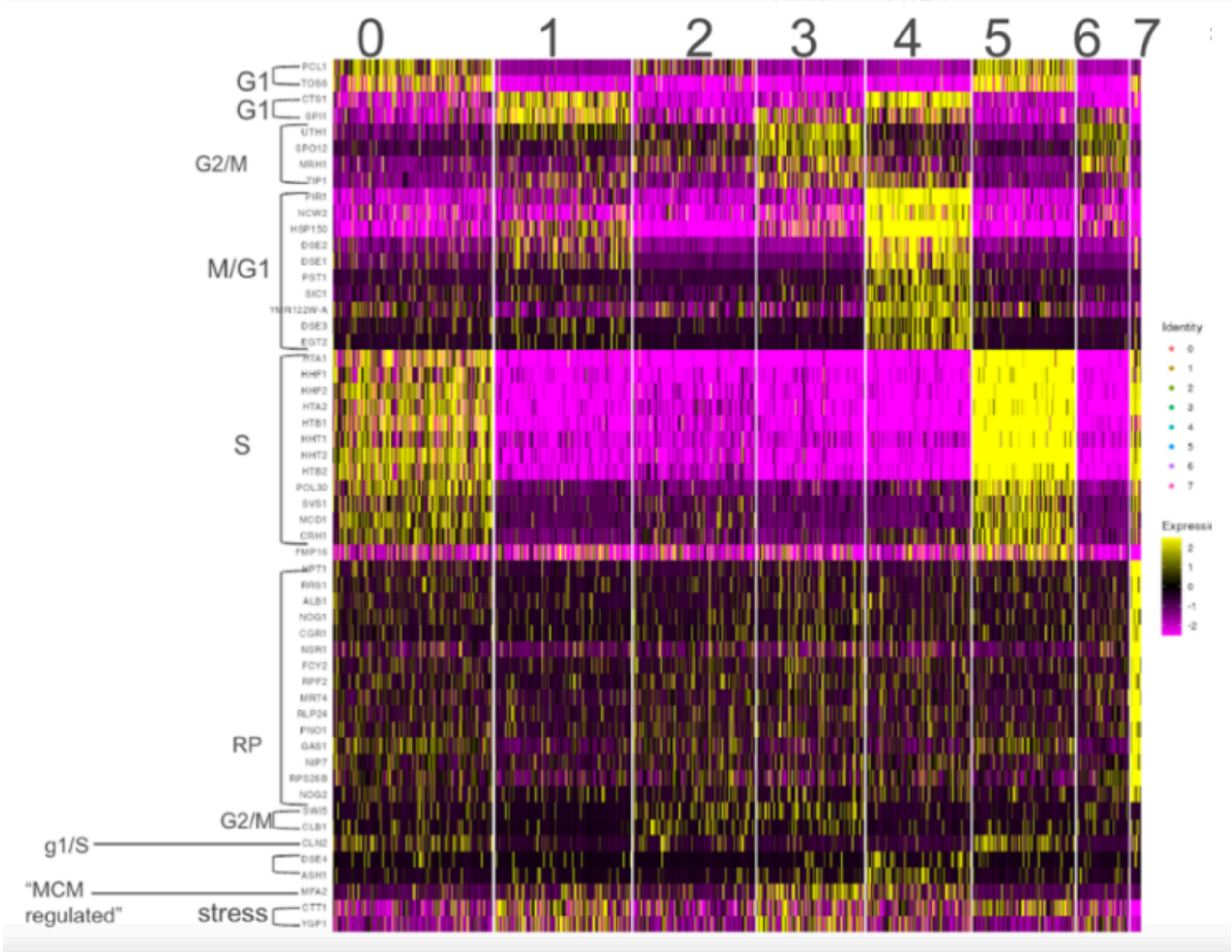


**Figure S26:** Clusters produced from BYxRM NaCl t30 at resolution 0.43. The stress cluster is no longer produced but the RP+S cluster persists. Other clusters can be assigned to a gross CC phase using expected marker expression.

We assign M/G1 to cluster 4 based on conventional cc markers such as PIR1. Unlike in the t0 condition, we do not see distinct clusters of M/G1 expression where mating genes are either on highly or nearly not measured at all so have only one ‘M/G1’ state in this dataset.

We assign G1/S using CLN2 and S phase markers that are much fainter in cluster 0 than in 5. We assign S as cluster 5 since the S band of genes is very highly expressed. We collapse cluster 7 with S since it has some S phase gene expression and is too small to conduct eQTL mapping in. Note that POL30 is very high in cluster 0 and not in cluster 5. Based on the expression in t0, where pol30 is high in G1/S and not in S, we will keep 0 and 5 distinct and assign G1/S to 0 and S to 5.

We assign G1 to cluster 1 based on cts1 expression and a lack of conflicting markers. We note a similar proportion of G1 cells in the BYxRM NaCl t0 sample and it stands to reason that a similar proportion of cells in a similar state could exist at t30 although we do not expect all CC markers to be interpreted exactly the same way so soon after a salt perturbation.

G2/M is assigned by using bands of G2/M markers expressed most clearly in 2, 3 and 6. Because these G2/M clusters have very distinct expression of mating genes, we left these as separate states for eqtl and state qtl mapping such that clusters 2 and 6 are collapsed as G2/M without mating and cluster 3 is assigned ‘G2/M + mating’.

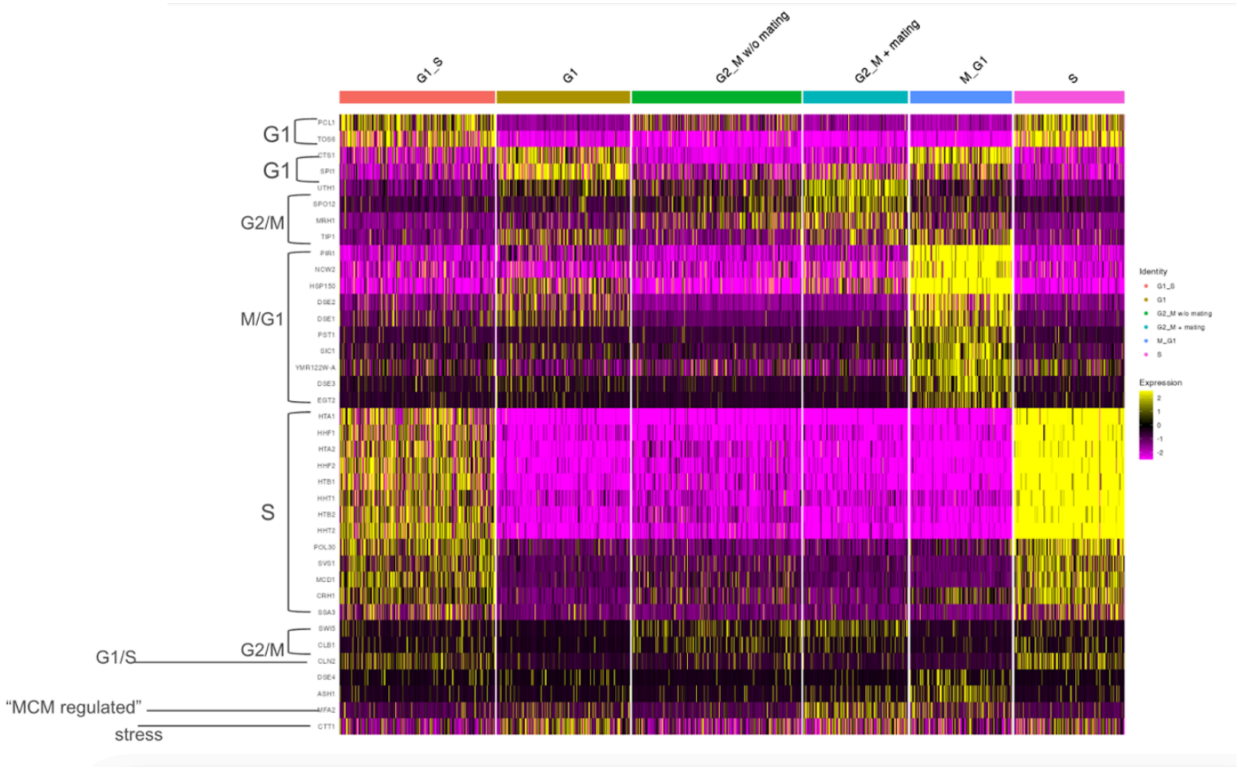


**Figure S27:** Clusters produced from BYxRM NaCl t30 at resolution 0.43 collapsed into default cc phases.

#### CBSxYJM NaCl t0

Processing of the CBSxYJM NaCl t0 sample was similar to that of BYxRM NaCl t0 until the PCA-based clusters were initially inspected using a heatmap. We noticed cluster 4 did not appear to have distinguishing expression of CC markers and expressed most genes at lower levels than nearly all other clusters. We removed this cluster and repeated sctransform, pca, and clustering

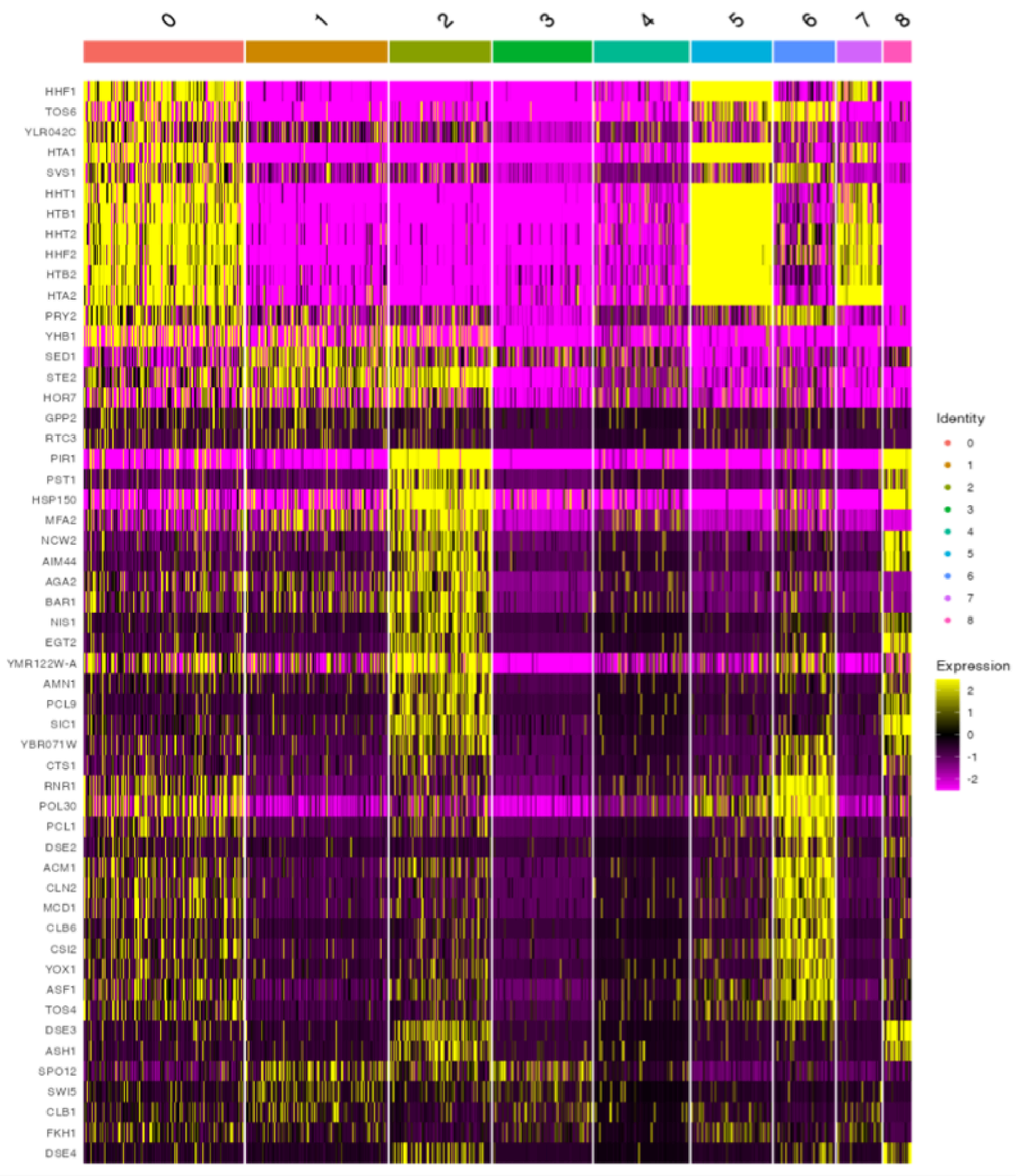


**Figure S28**. CBSxYJM NaCl t0 PCA-based clusters prior to removal of cluster 4.

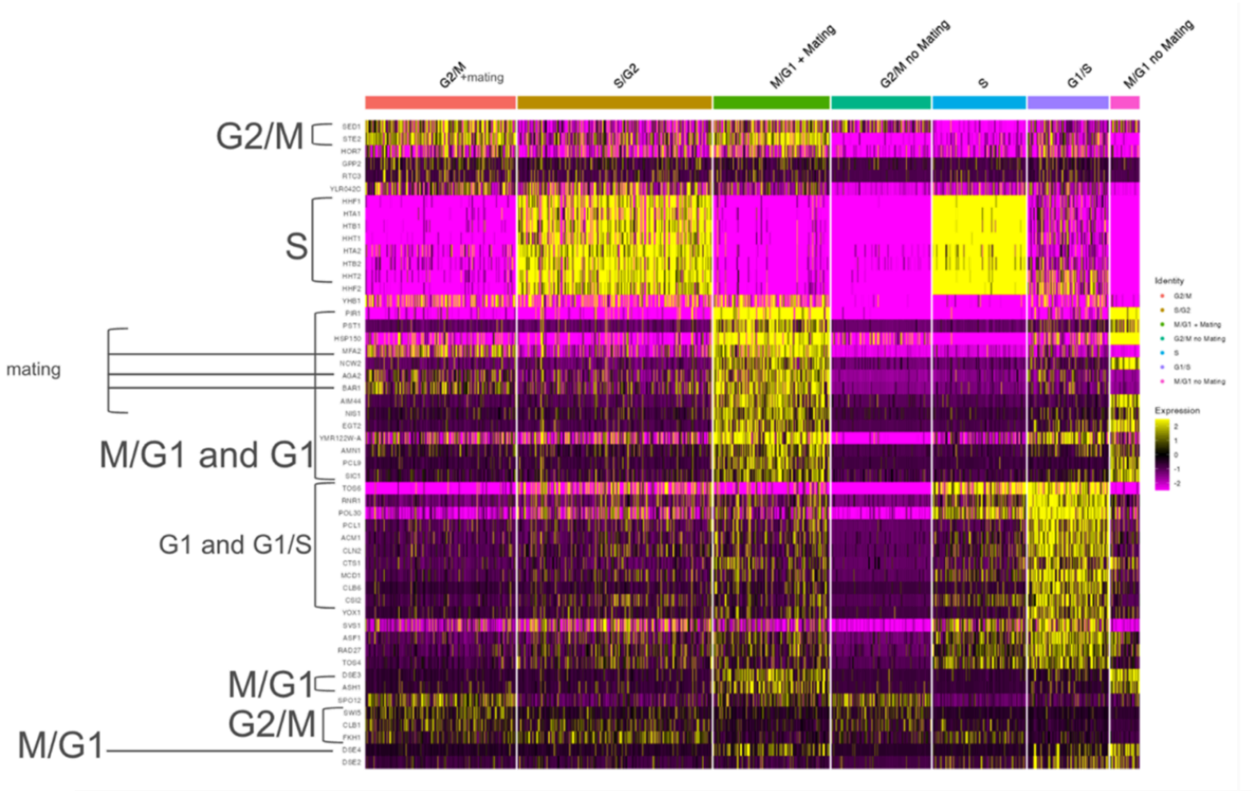


**Figure S29.** Moderate resolution clustering in 12 dim pc space of CBSxYJM NaCl t0 dataset after removal of cluster 4.

The resulting clusters resembled those from the BYxRM t0 experiment in most regards with one difference being that there appeared to be no clusters like cluster 4 in BYxRM NaCl t0 (pre collapsing clusters) that had a distinct G1 character (distinct CTS1 and DSE2 expression, for example, which is generally lower in the CBSxYJM experiment but is much higher in clusters 2 and 5 (above, “M/G1 + mating” and “G1/S”). Bands of M/G1 expression can be seen in clusters 2 and 7 with the difference between these being mating gene expression. Similarly, clusters 0 and 3 share characteristic G2/M expression but differ in terms of mating gene expression. At much higher resolution, we did not see evidence of a small population of cells that could be designated as G1 based on the criteria used for the BYxRM t0 sample.

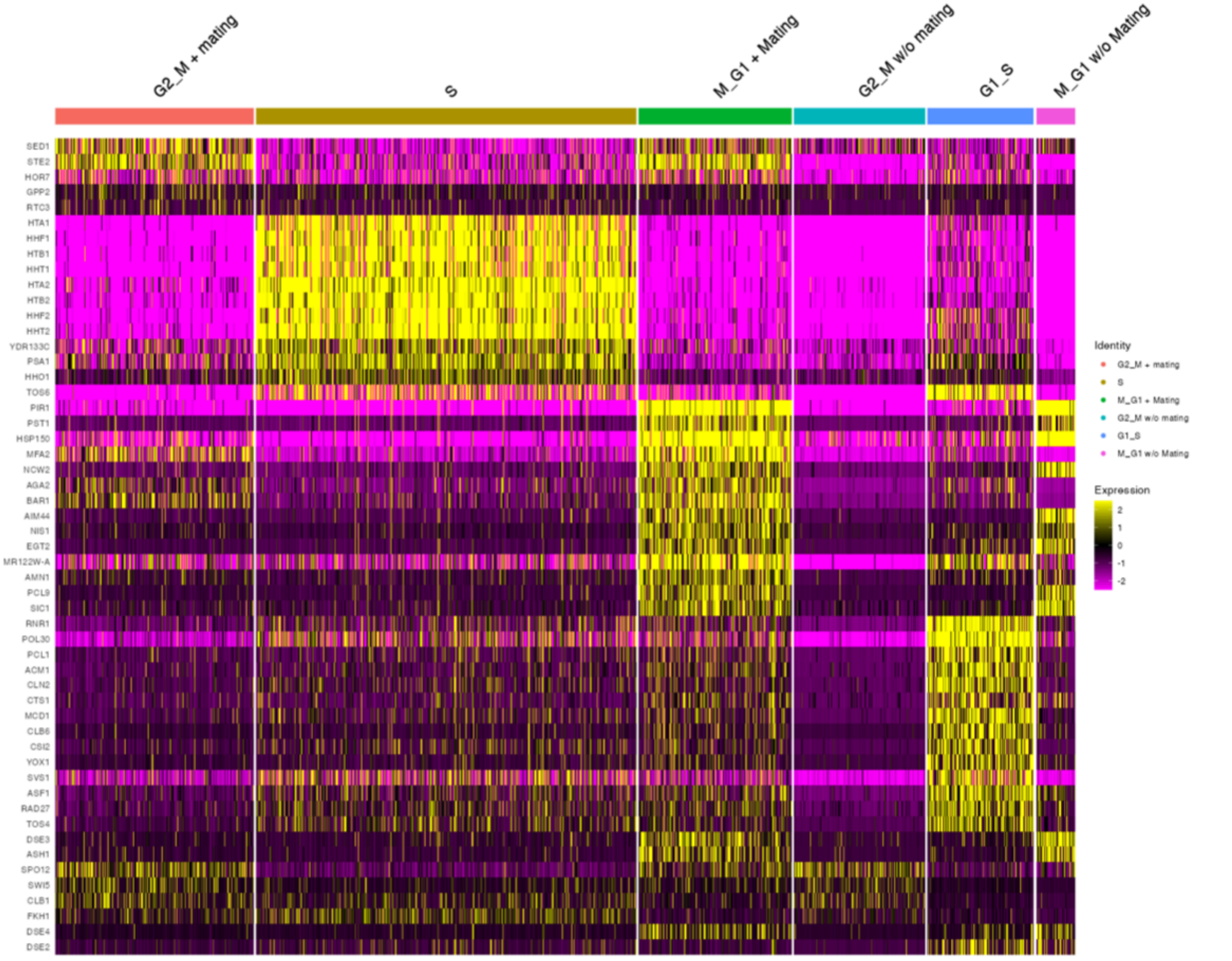


**Figure S30.** Cell cycle labels applied to medium resolution clusters were collapsed to the same states used in BYxRM NaCl t0 without an explicit 'G1'.

**CBSxYJM NaCl t30**

High resolution clustering in 12 dim pca space after removal of alphas and variance-stabilization with sctransform shows a similar 'unknown' set of cells with unusual/indistinct and muted expression profiles as was seen and removed from the CBSxYJM NaCl t0 sample. We removed this 'unknown' cluster and repeated sctransform as well as PCA using the Spellman et al. CC regulated genes. We found a large cluster distinguished by stress response gene expression that persists at various clustering resolutions. In contrast, the BYxRM t30 sample has a similar cluster at very high resolution but at lower resolution it does not persist and at high resolution it is very small. There is also a smaller cluster in this sample with very strong ribosomal protein gene expression and moderate s phase marker expression. We chose to collapse this cluster with s phase manually based on the marker expression but note that it does have a very distinct character.

Unlike the RP + S phase cluster, the cluster with distinct stress response gene expression is much larger. There is faint expression of cc markers for several phases and it did not seem obvious how to place it within the conventional cc labels we have used in mid-log ypd/t0 conditions. We note that the stress cluster in BYxRM has less expression of cc markers at all vs. confusing expression of multiple cc markers at once. We chose to leave it and annotate it as ‘stress’ unlike our decision to use a low enough clustering resolution for the BYxRM t30 data that the (much smaller) analogous stress cluster is partitioned into larger clusters. In both cases, we manually collapse the RP +s phase cluster with S phase (in BYxRM t30, even at low res it shows up as a small number of cells with distinct RP expression).

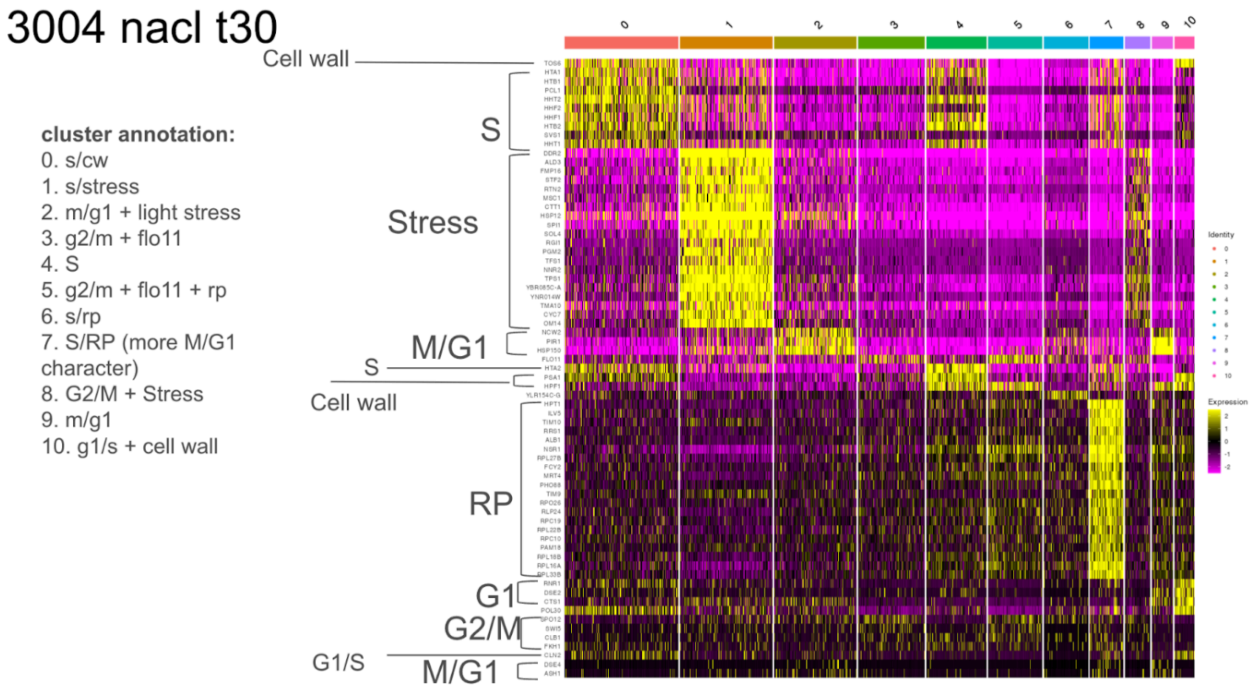


**Figure S31.** High-resolution clustering of CBSxYJM NaCl t30 prior to collapsing clusters.

Because the stress gene expression was much higher in 3005 NaCl t30 vs BYxRM NaCl t30, we decided to include this as a distinct state. Cluster 1 has the most distinct expression of the band of stress genes (CTT1, for example) but cluster 8 also has distinct, though much fainter, expression of these genes along with faint expression of G2/M markers. We collapsed clusters 1 and 8 into a general stress cluster.

To assign G1, we note that CTS1 is most distinct in cluster 10 and that CLN2 is also expressed (a G1/S marker that may show up in G1 and persist through S phase). There are not conflicting markers expressed in cluster 10 so we label it G1. Interestingly, this state does not appear in the CBSxYJM t0 sample but seems to be represented by a small number of cells in the stimulated condition. In comparison, a distinct G1 state appears in both the t0 and t30 stimulated NaCl BYxRM samples.

To assign M/G1, we use expression of m/g1 markers such as PIR1 to assign clusters 2 and 9 as m/g1 but note that cluster 9 has more rp gene expression (not as strongly as in the S + RP cluster/cluster 7). Unlike in the BYxRM NaCl t30 sample, we saw clusters of M/G1 expression with and without mating genes. Although we did not see this in our very high-resolution clustering in BYxRM NaCl t30, we decided to assign m/g1 with and without mating gene expression in CBSxYJM. It is possible that at much higher resolution

To assign G1/S and S, we used the expression of CLN2 as well as canonical s phase expression. In no cluster was s phase expression as extremely distinct and high as during mid log but both clusters 0 and 4 show clearly high s phase gene expression. Clusters 1 and 7 do also have low expression of classic s phase genes but this is relatively low in cluster 1 and very moderate in 7. The concurrent strong expression of s phase genes and RP genes in cluster 7 speaks to its distinctness but it could reasonably be collapsed into S phase or G2/M based on low expression of markers for both phases. Because G1/S marker CLN2 expression is fairly high here vs the clusters we assign as G2/M (3 and 5), it made most sense to assign cluster 7 as S phase (probably the tail end of s or a transition to G2/M?). POL30 and CTS1 are moderately high in cluster 0 but less so in the other clusters with s phase expression so we assign cluster 0 G1/S and note that it has lower expression of s phase markers than the clusters we do call s but it also has more g1 character than those clusters (looking at the pol30 line makes that point). We collapse clusters 4 and 7 as S phase.

To assign G2/M phase, we noted that clusters 3 and 5 have moderate expression of G2/M markers (CLB1, SPO10, SWI5, etc.) and lacked concurrent expression of markers from other phases. We note that several clusters have low levels of G2/M marker expression in this experiment but that G1 marker POL30 is very low in these clusters and that the G2/M markers shown in the above heatmap are very low in the cluster being classified as G1. Cluster 6 has expression of SPO12, a G2/M marker that is higher than in some other clusters/states (for example, M/G1, G1, and G1/S have virtually no expression of this gene) so we assigned it a label of G2/M but note that it does have distinct expression of HPF1 relative to all other cells in this sample.

We collapsed these clusters into the standard CC phases from the mid-log/t0 condition but kept the ‘stress’ cluster/cluster 1 instead of attempting to collapse it into another phase assignment or to cluster everything at so low a resolution that it would disappear. Although we can not use the stress cluster in the BYxRM NaCl t30 dataset for mapping because this subset of cells is too small/won’t afford necessary power, we can do so here.

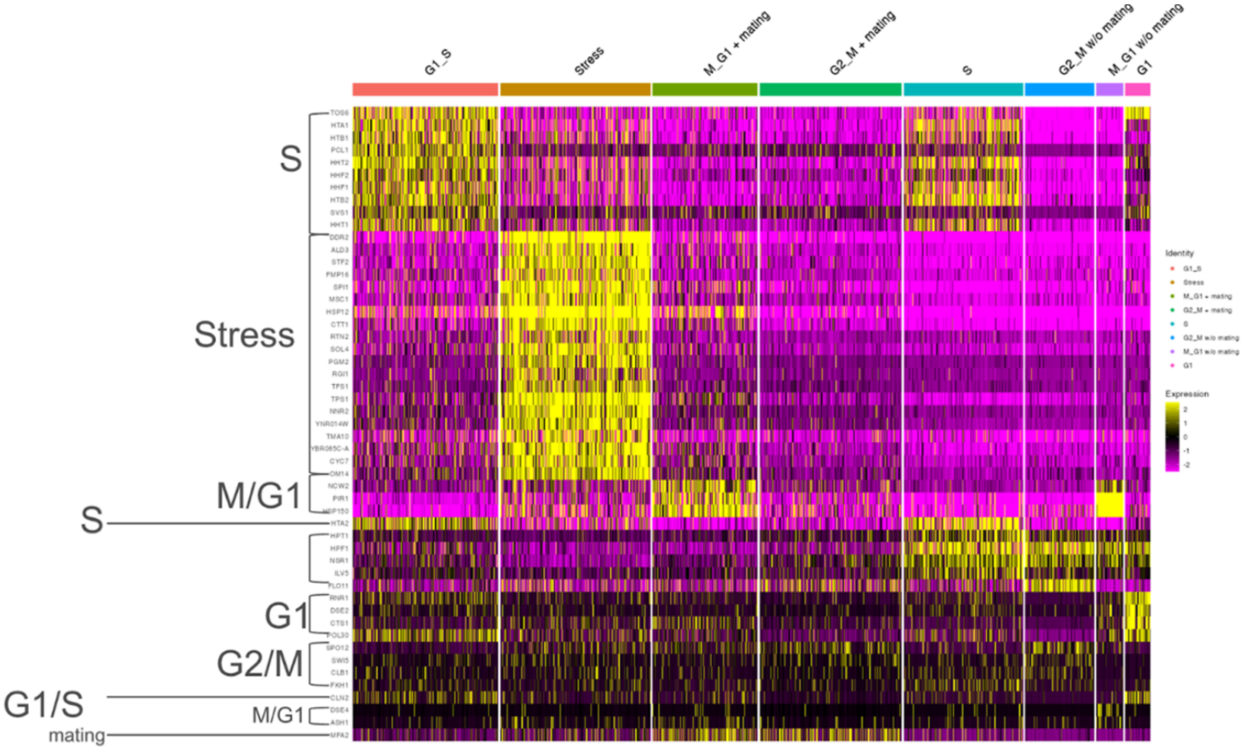


**Figure S32.** CBSxYJM NaCl t30 collapsed state labels.

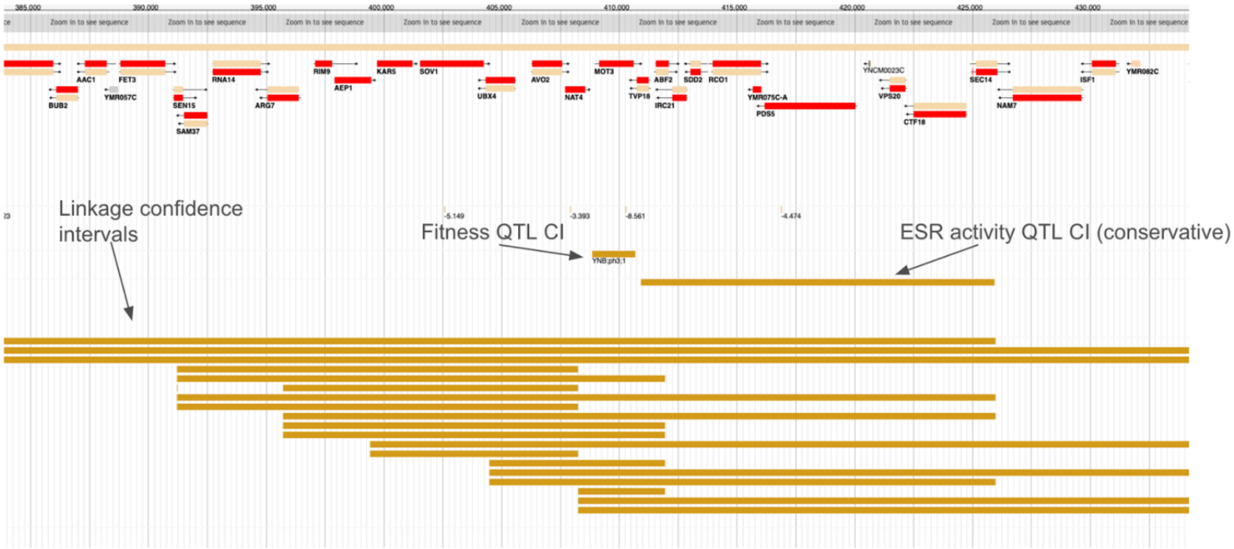


**Figure S33.** Example of an eQTL hotspot hypothesized to be driven by variation in the MOT3 TF overlapping other QTL.

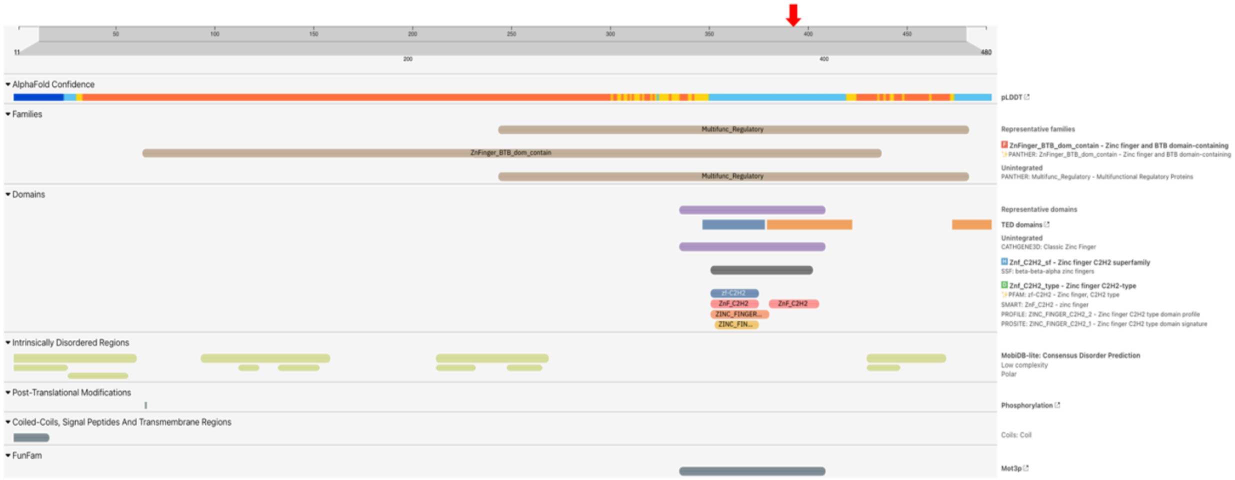


**Figure S34.** Location in MOT3 protein where a hypothesized causal missense variant is situated. The location of the variant is denoted by a red arrow at the top of the figure.
